# Overlooked features lead to divergent neurobiological interpretations of brain-based machine learning biomarkers

**DOI:** 10.1101/2025.03.12.642878

**Authors:** Brendan D. Adkinson, Matthew Rosenblatt, Huili Sun, Javid Dadashkarimi, Link Tejavibulya, Corey Horien, Margaret L. Westwater, Raimundo X. Rodriguez, Stephanie Noble, Dustin Scheinost

**Author notes:** Corresponding author: Brendan Adkinson, Magnetic Resonance Research Center, 300 Cedar Street, New Haven, CT, 06519.

## Abstract

A central objective in human neuroimaging is to understand the neurobiology underlying cognition and mental health. Machine learning models trained on brain connectivity data are increasingly used as tools for predicting behavioral phenotypes 1,2, enhancing precision medicine 3,4, and improving generalizability compared to traditional MRI studies 5. However, the high dimensionality of brain connectivity data makes model interpretation challenging 6. Prevailing practices within the field rely on sparsely selected brain connectivity features, implicitly interpreting identified feature networks as uniquely representative of a given phenotype while overlooking others. Here, we show that commonly overlooked brain connectivity features can achieve similar prediction accuracies while yielding markedly different neurobiological interpretations. Using four large-scale neuroimaging datasets spanning over 12,000 participants and 13 outcomes, we demonstrate that this phenomenon is widespread across cognitive, developmental, and psychiatric phenotypes. It extends to both functional connectivity (fMRI) and structural (DTI) connectomes and remains evident even in external validation. These findings suggest that common practices may lead to feature overinterpretation and a misrepresentation of the neurobiological bases of brain-behavior associations. Such interpretations present only the "tip of the iceberg" when certain disregarded features may be just as meaningful, potentially contributing to ongoing issues surrounding reproducibility within the field. More broadly, our results point to the possibility that multiple neurobiologically distinct models may exist for the same phenotype, with implications for identifying meaningful subtypes within clinical and research populations.

## Introduction

A central goal of human neuroimaging is to uncover the neurobiological underpinnings of cognition and mental health. Aligned with this objective, emerging machine-learning approaches predict individual differences in behavioral phenotypes from brain structure and function ^1,2,7,8^. These ‘brain-behavior’ predictive models reduce overfitting and provide more robust assessments of associations relative to traditional magnetic resonance imaging (MRI) studies by separating datasets into training and test samples ^5,9–11^. They also offer opportunities to advance precision medicine strategies, ranging from illness subtyping to predicting individual treatment responses ^3,4,12^.

Accordingly, substantial efforts have been dedicated to improving model accuracy ^13–15^, trustworthiness ^16^, fairness ^17,18^, reproducibility ^19–24^, and generalizability ^25–28^.

Neurobiological interpretability—the degree to which results can be mapped to known neural circuits and processes—is another key metric of model utility ^29–31^. An easily interpretable model can uncover the neural circuits responsible for behavior, even with modest prediction performance ^32^. However, several challenges exist for interpreting machine learning models based on neuroimaging data ^6,33,34^. To obtain more manageable representations of high dimensional data, the field commonly employs univariate feature selection ^35,36^. Here, brain features are associated one-by-one with the target phenotype. Those with the strongest univariate effect sizes are selected for modeling, while all others are discarded. Although feature selection can reduce training time, improve predictions, and simplify interpretability, it may overlook weaker, neurobiologically meaningful signals.

Traditionally, selected features are interpreted as the neurobiology underlying individual variation in the predicted phenotype. For instance, models are often named after the phenotype they predict (e.g., “working memory network” or “depression network”). By treating selected features as definitive, these practices give the impression that a model uniquely represents that phenotype’s neural correlates.

Implicitly, unselected features are overlooked as less important—or, in some cases, inconsequential—to model performance and neurobiology.

As a result, feature selection may risk misrepresenting the neurobiology underlying individual differences in behavior. Associations between the brain and real-world phenotypes are likely best characterized by widely distributed neural circuits with small effect sizes ^23,37,38^. Conventional univariate approaches, including univariate feature selection, may permit only the most robust and straightforward associations—the tip of the iceberg—to survive ^39^. Yet, recent work demonstrates that features with weaker univariate effect sizes can often be combined to yield comparable prediction accuracy to more strongly associated features ^12^. Feature selection may thus oversimplify the complex neurobiology subserving behavior. If features discarded during selection support meaningful predictions, then restricting interpretation to top-ranked features truncates neurobiological complexity. Despite its widespread use, how univariate feature selection balances the tradeoff between simplification for optimizing modeling and oversimplification that misrepresents true neurobiology remains understudied.

Here, we test whether unselected features are meaningful for prediction and neurobiological interpretation. Our analyses span 12,200 participants across four large-scale neuroimaging datasets, both functional and structural connectivity, and 13 outcomes, including age, sex, cognitive abilities, developmental measures, and psychiatric phenotypes. We use an original prediction paradigm wherein connectome features are divided into non-overlapping subsets (i.e., an edge can only be in a single subset) based on their association with a target phenotype. We show that multiple subsets overlooked by feature selection can yield significant prediction accuracies but diverging biological interpretations. These results suggest that focusing on the “top” features may potentially paint an incomplete picture of the underlying neurobiology and reinforce that subtle brain-wide signals should not be ignored. They also suggest that multiple neurobiologically-distinct models may exist for a given phenotype, which could have important implications for identifying meaningful subtypes within clinical or research populations.

## Results

### Overview

We used four developmental neuroimaging datasets: the Healthy Brain Network (HBN, n=1110) dataset ^40^, the Adolescent Brain Cognitive Development (ABCD, n=9371) Study ^41^, the Human Connectome Project Development (HCPD, n=428) dataset ^42^, and the Philadelphia Neurodevelopmental Cohort (PNC, n =1291) dataset ^43^. Details about the datasets are presented in Table S1 and in the Methods section; see also Adkinson et al. (2024) ^25^ for further context. Our first set of analyses focused on connectome-based predictive modeling, which explicitly relies on univariate feature selection. We later present complementary results using ridge regression (which can handle correlated features without feature selection) with and without feature selection.

We modified connectome-based predictive modeling ^44^ such that connectome features were divided into ten non-overlapping deciles based on the strength of their association with a target phenotype (Figure 1A). Specifically, to prevent data leakage, we computed the Pearson correlation coefficient between every connectivity edge and the target phenotype in the training set. Features were then ranked in descending order based on the absolute value of their correlation coefficients. Following ranking, the complete set of connectivity features was partitioned into ten non-overlapping deciles, with the first decile comprising the top 10% of features (*i.e.*, those most likely included in established neuroimaging predictive modeling pipelines) and the last decile comprising the bottom 10% of features. Each decile was subsequently used to predict the phenotype of interest in the respective test set. Models were trained using 100 iterations of 10-fold cross-validation. Model performance was evaluated with Pearson’s correlation (r), representing the correspondence between predicted and actual behavioral scores, along with the cross-validation coefficient of determination (q^2^) and mean square error (MSE). Significance was assessed using permutation testing with 1000 iterations of randomly shuffled behavioral data labels. As in previous work, the Haufe transform was not applied to the CPM models^33,45^.

**Figure 1.**
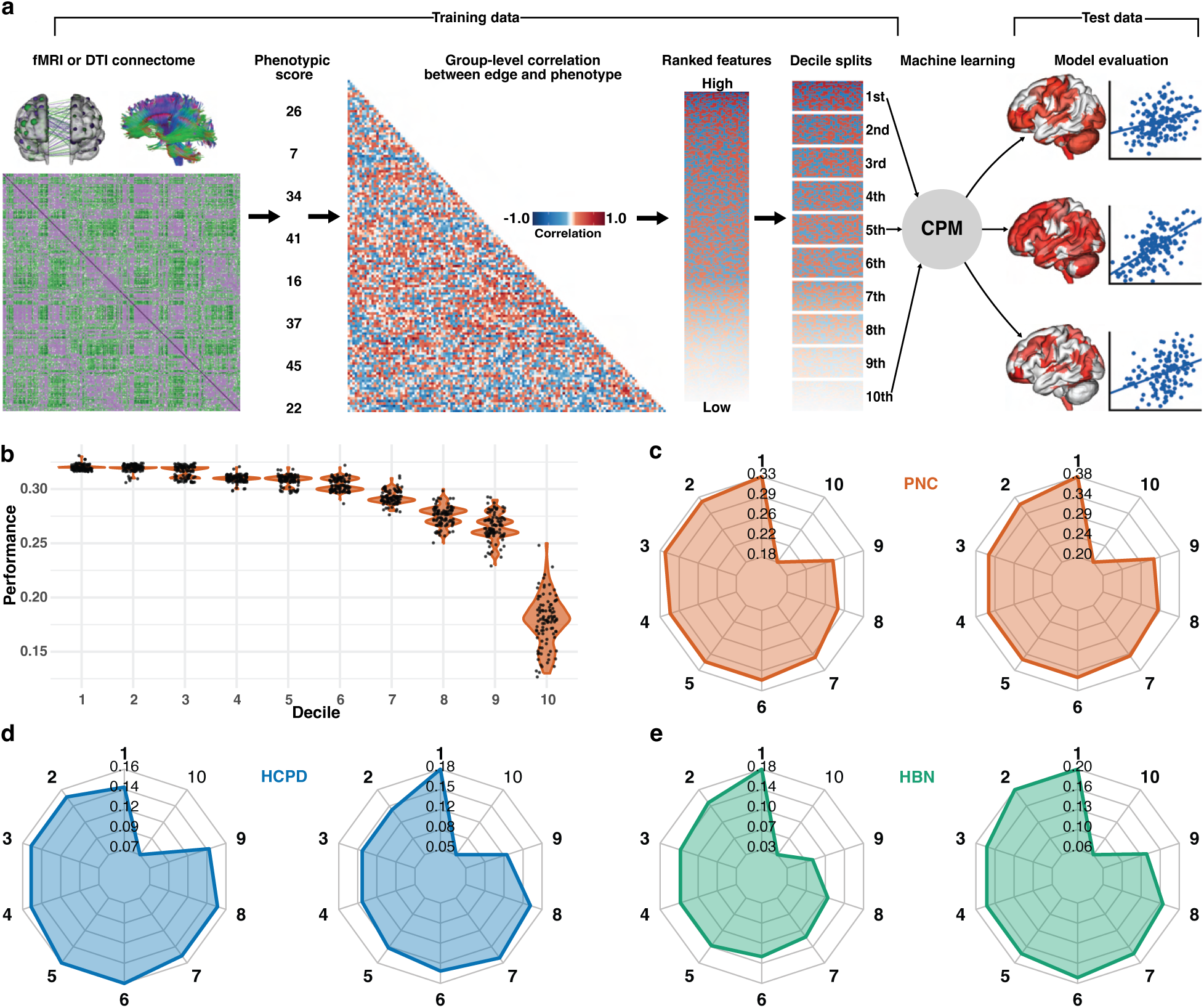
Connectome-based predictive modeling (CPM) across non-overlapping decile-ranked brain connectivity features. (A) Workflow illustrating the decile-based connectome-based predictive modeling (CPM) pipeline, including initial correlation of connectivity features with phenotypic outcome, ranking features based on group-level correlations between edges and phenotype, splitting features into deciles, and evaluating each decile-based model. (B) Violin plot showing the predictive performance of models trained on each decile of features within the PNC dataset for executive function. (C-E) Radar plots depicting predictive performance across deciles for PNC executive function (C, left), PNC language abilities (C, right), HCPD executive function (D, left), HCPD language abilities (D, right), HBN executive function (E, left), and HBN language abilities (E, right). Bolded decile numbers indicate significant predictions.

### Prediction accuracy is not exclusive to top-ranked features

We generated language abilities and executive function models in the PNC, HBN, and HCPD datasets using our modified CPM paradigm. As in previous work ^25^, we used principal component analysis (PCA) to combine individual cognitive measures into latent variables of executive function and language abilities. Models constructed from the first through ninth feature deciles successfully predicted executive function and language abilities across PNC, HBN, and HCPD (Figure 1, Figures S1-3). Lower-ranked features commonly overlooked during model building continued to demonstrate significant prediction performance. For example, for PNC executive function, the first decile achieved a prediction performance of *r*=0.33 (p=0.001, q2=0.09, MSE=1.24), while the second (*r*=0.32, p=0.001, q2=0.08, MSE=1.24) through sixth (*r*=0.31, p=0.001, q2=0.07, MSE=1.25) deciles achieved significant predictions. Notably, while these models perform similarly, the underlying edge features are non-overlapping, in that a given edge could only be in one decile. Additionally, the first decile did not always exhibit the best prediction performance. For HCPD executive function, the fifth decile (*r*=0.16, p=0.001, q2=-0.07, MSE=2.08) numerically outperformed deciles with features ranked as more highly informative (e.g., first decile *r*=0.14, p=0.002, q2=-0.10, MSE=2.14). Overall, the first decile showed no significant differences in prediction performance compared to the 2nd, 5th, 6th, and 7th decile (Supplementary Table S2). Performances for all deciles are presented in Supplementary Table S3.

### Overlooked feature sets also pass external validation

As external validation is the gold standard for model evaluation ^24^, we applied our decile-based models to three independent datasets. Despite the absence of overlapping features between deciles, deciles beyond the first demonstrated successful external validation for executive function and language abilities across all 12 cross-dataset prediction scenarios (Figure 2, Tables S4, S5, S6). For example, PNC executive function models tested in HCPD had no significant differences in external validation between deciles 1 through 9. In other words, the PNC executive function models from the 9^th^ decile (*r*=0.13, *p*=0.004) generalized to HCPD just as well as the 1st decile (*r*=0.14, *p*=0.002). Similar trends were observed for other models. Models tested in PNC (*r*’s generally >0.25, Figure 2) and HCPD (*r*’s generally >0.10) outperformed those tested in HBN (*r*’s generally <0.10).

**Figure 2.**
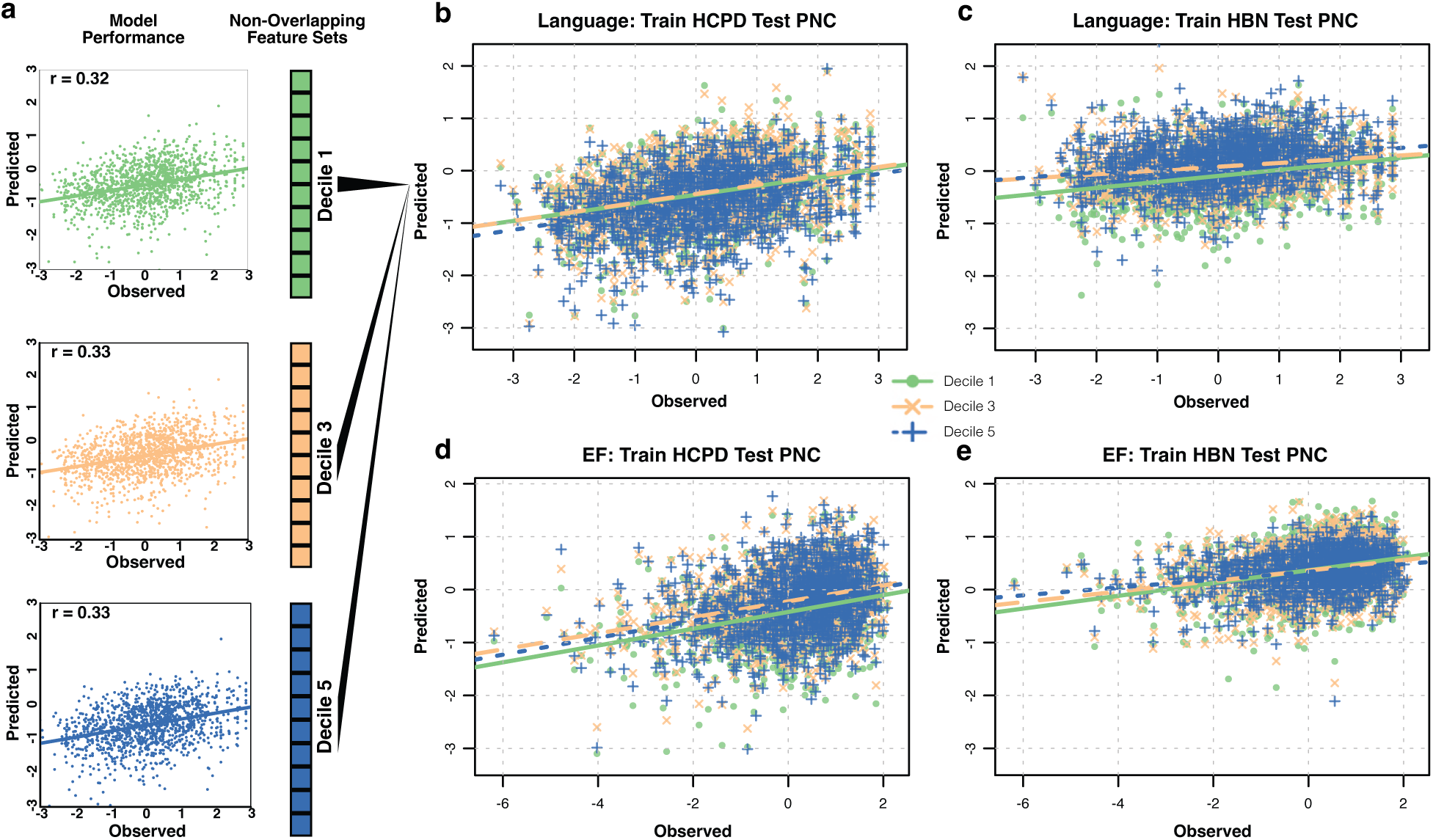
Multiple non-overlapping feature sets are generalizable to external datasets. (A) Scatter plots depicting observed versus predicted values for language abilities predictions when training in HCPD and testing in PNC using 1st (green), 3rd (blue), and 5th (purple) decile feature subsets. (B-E) Scatter plots for 1st (green), 3rd (blue), and 5th (purple) decile feature subsets overlaid to enable direct comparison of predictive accuracy across deciles. (B) Training a language abilities model in HCPD and testing in PNC. (C) Training a language abilities model in HBN and testing in PNC. (D) Training an executive function model in HCPD and testing in PNC. (E) Training an executive function model in HBN and testing in PNC.

### Overlooked features yield diverging neurobiological interpretations

Next, because models performed similarly across non-overlapping feature subsets, we sought to determine if similar brain features were implicated in each decile-based model by comparing within-and between-network functional connectivity. The canonical networks underlying predictions differed between deciles for both executive function and language abilities (Figure 3, Figure S4). For example, for PNC executive function, connectivity between the visual association and frontal-parietal networks was prominent in decile 1 but less informative for deciles 2 through 5 (Figure 3B). Networks from different deciles showed weak to moderate explained variance (*average* = 13%, Figure S5), with only one network pair explaining greater than 50% variance. Explained variance between networks from neighboring deciles was greater than explained variance between those from non-neighboring deciles. For PNC executive function, the explained variances were 0.5% between decile 1 and decile 3, 3% between decile 1 and 4, and 4.5% between decile 1 and 5 (Figure S5). That networks became increasingly dissimilar with further deciles was consistent across datasets and phenotypes. Trends were also similar when shifting from a network-based comparison to a node-based approach, evaluating features based on the sum of model contributions for each of the 268 nodes (Figure 3C).

**Figure 3.**
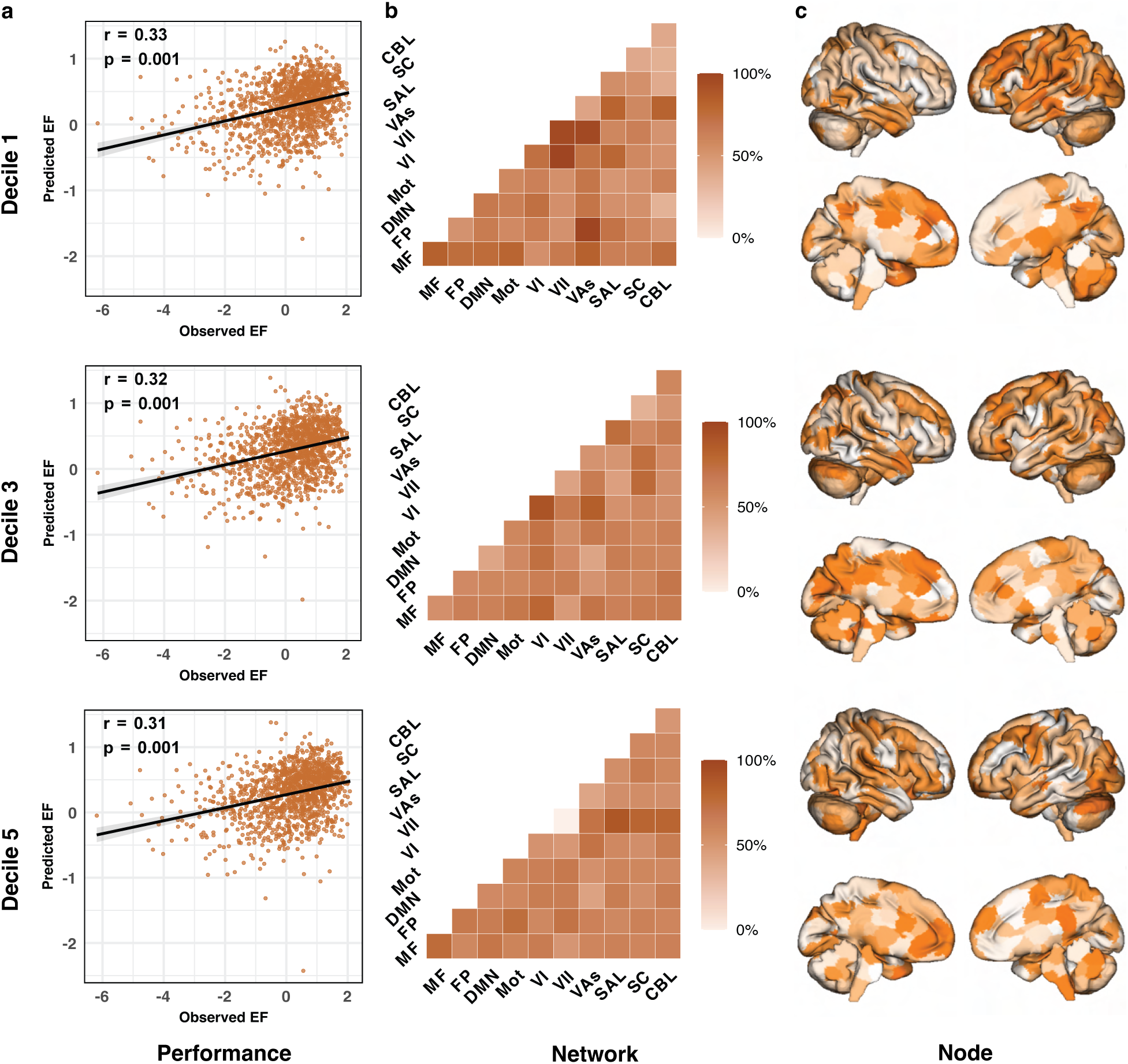
Overlooked feature sets offer similar prediction performance but unique neuroanatomical contributions. (A) Scatter plots depicting model performances across deciles as measured by the Pearson correlation between observed and predicted values. Gray bands show the 95% confidence interval. One-sided p values are reported. (B) Canonical network contributions to PNC executive function predictions. Contributions represent the number of selected features averaged across folds (i.e., proportion of folds in which each edge was selected in the median-performing model) grouped by canonical functional network pairs. Diagonal cells represent contributions of edges within a single network; off-diagonal cells represent contributions of edges between networks. Values were normalized by the number of edges in each network group. Darker colors indicate higher relative contribution. Network Labels: MF, medial frontal; FP, frontoparietal; DMN, default mode; Mot, motor cortex; VI, visual A; VII, visual B; VAs, visual association; SAL, salience; SC, subcortical; CBL, cerebellum. (C) Node-level contributions to PNC executive function predictions. Node contributions show the sum of all edgewise ridge regression coefficients averaged across folds.

The selected features were more frequently present across folds in the higher deciles than in the lower deciles (Table S7). On average across all models, relative to the first decile, 84.64% of second-decile edges, 38.82% of third-decile edges, and 22.19% of fifth-decile edges were present (i.e., selected) in at least five of the ten cross-validation folds. To account for this variation, the proportion of folds in which each edge was selected for CPM models was used for all interpretability analyses (see Methods). Analyses were repeated using edges selected consistently across at least five folds, yielding similar results (Figure S6).

### Utility of overlooked features generalizes to psychiatric, developmental, and demographic phenotypic domains

Next, we examined whether overlooked features have relevance in domains beyond language and executive function. First, predictions were performed within a diverse selection of developmental and psychiatric phenotypes in the HBN dataset (n=747 participants, see Methods). For every phenotype, deciles beyond the first retained predictive utility (Figure 4A). Notably, for the Social Communication Questionnaire ^46^, the second through seventh deciles achieved numerically greater prediction performance (*r*’s=0.15-0.16, MSE’s=19.61-19.73, q2’s=-0.02) than the first decile (*r*=0.14, q2=-0.03, MSE=19.80). Predictive models of cognitive, developmental, and psychiatric phenotypes generally exhibit low effect sizes. As such, we also tested whether our findings extend to predictions of larger effect size phenotypes by predicting age and sex in the PNC, HCPD, and HBN. Once again, traditionally overlooked feature sets retained significant predictive capabilities for sex and age in all three datasets (Figure 4B, C).

**Figure 4.**
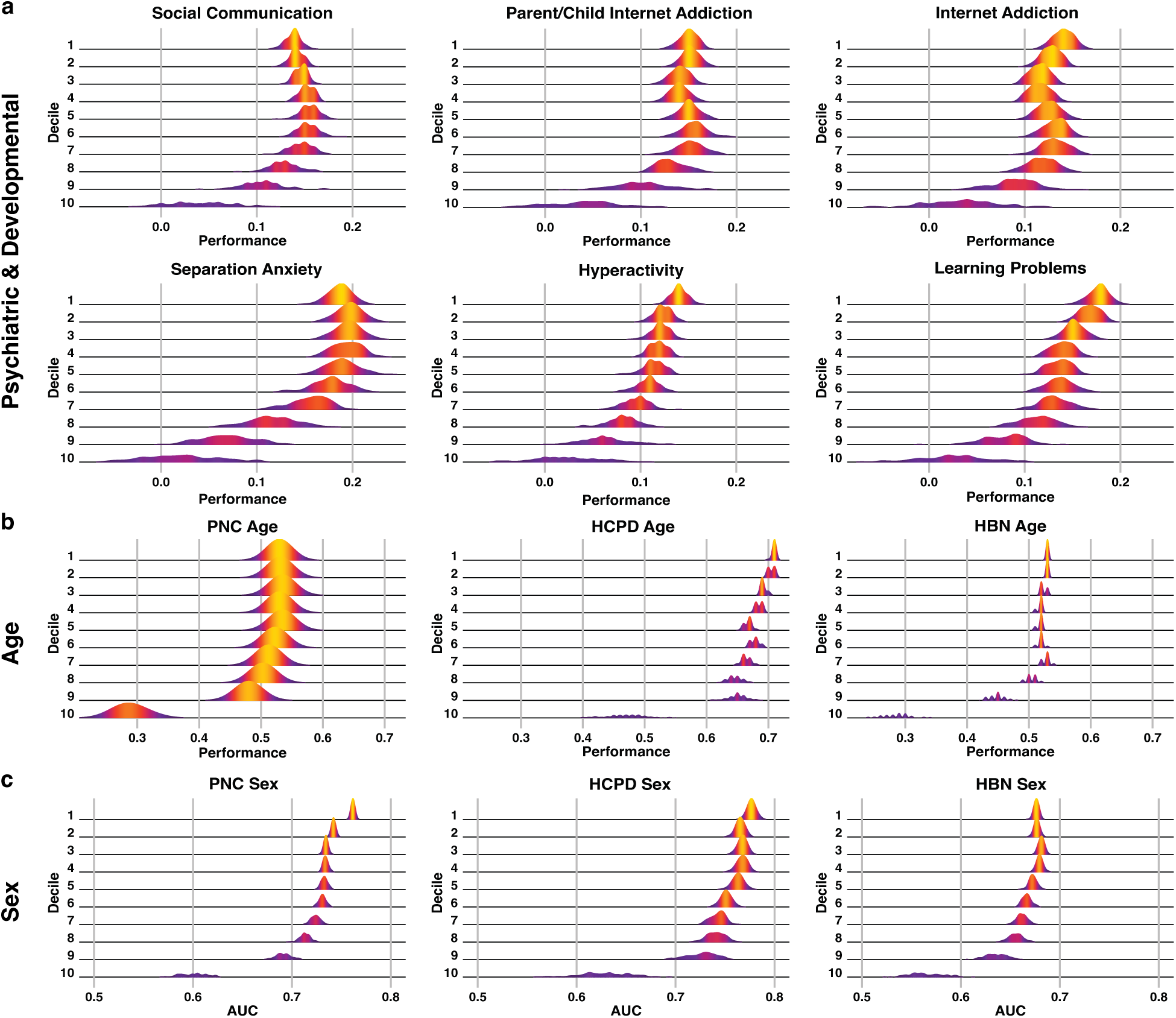
Decile-based predictive modeling performance across psychiatric, developmental, and demographic phenotypes in HBN. (A) Ridgeline plots showing the distribution of prediction performances (Pearson’s r) across 100 iterations for the Social Communication Questionnaire (SCQ), Strengths and Difficulties Questionnaire (SDQ) Hyperactivity, Screen for Anxiety Related Disorders (SCARED) Separation Anxiety, Parent-Child Internet Addiction Test (PCIAT), Internet Addiction Test (IAT), and Conners 3 Self-Report (C3SR) Learning Problems scores in HBN. (B) Predictive performance of age for PNC, HCPD, and HBN datasets. (C) Predictive performance of sex in the PNC, HCPD, and HBN datasets.

### Overlooked features with ridge regression

Next, we repeated several analyses using ridge regression. Ridge regression can handle high-dimensional data without the need for feature selection ^47^. Though, feature selection is still commonly employed with ridge regression. Ridge regression was performed both without feature selection and with using an analogous decile-based feature selection approach as the previous results. For biological interpretation, the Haufe transform^48^ was applied to the model weights.

Feature selection increased prediction performance and decreased training time for ridge regression. Ridge regression models without feature selection performed significantly worse than the first decile models (p=0.023) and similar to the second (p=0.055) and third decile models (p=0.660, Tables S8 and S9). Models from all other deciles performed worse. Additionally, the decile-based models were significantly quicker to train (p<0.01). These results highlight the benefit of feature selection, even when methods do not explicitly rely on it.

Similar to CPM, models beyond the first decile exhibited significant predictions (Figure 5A, p<0.05). Though, unlike CPM, prediction performance for the subsequent deciles were worse than previous deciles. For example, models from the first decile outperformed models from the second decile (p=0.048). Models based on non-overlapping deciles exhibited complementary connectivity patterns from each other and models without feature selection. Decile-based models typically explained less than 40% (at the network level) and 43% (at the node level) of the variance of the models without feature selection (Figures 5B). Only three decile-based models explained greater than 50% of the models without feature selection. In two-thirds of the comparisons, the most similar models to those without feature selection were not from the top decile model (Figure S7). In other words, models built on all features are (on average) more similar to lower decile models.

**Figure 5.**
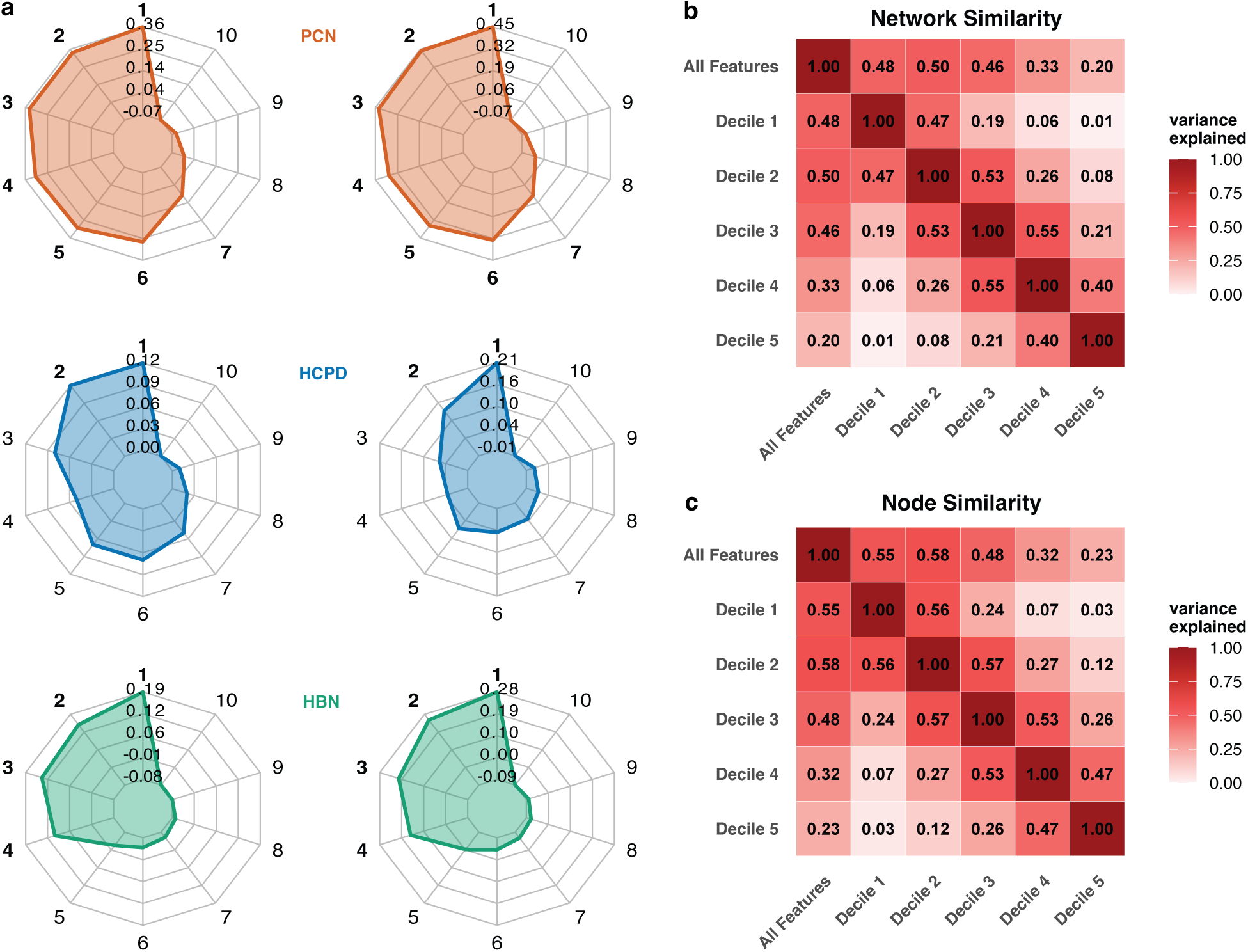
Decile-based model performance using ridge regression. (A) Radar plots depicting executive function (left) and language abilities (right) model performance across deciles for PNC (orange), HCPD (blue), and HBN (green). Bolded decile numbers indicate prediction performance r≥0.10. (B) Pairwise explained variance between models at the network level averaged across datasets. (C) Pairwise explained variance between models at the node level averaged across datasets. Higher explained variance reflects greater similarity in features across deciles. Supplementary Figure S7 presents results for individual datasets and phenotypes.

### Replication using partial correlation

fMRI features are known to have a high degree of spatial and temporal autocorrelations ^49,50^. To isolate direct connections by removing shared variance between nodes, we repeated our analyses using functional connectivity matrices generated with partial correlation instead of Pearson’s correlation in the PNC dataset. Partial correlation connectome results resembled those yielded by diffusion-based connectomes in that performances decayed more monotonically relative to the standard PNC functional connectomes (Supplementary Tables S10, S11).

### Overlooked features are also useful in other imaging modalities

We utilized diffusion weighted imaging data from the ABCD study to test whether the predictive utility of lower deciles is conserved across imaging modalities. Structural connectomes were generated based on each fiber’s average quantitative anisotropy value connecting two end regions (see Methods). We created functional and structural models for the 6271 ABCD participants with both fMRI and diffusion data to predict NIH Toolbox age-corrected fluid, crystallized, and composite intelligence scores.

Like functional models, structural models retained considerable predictive abilities throughout lower decile feature sets (Figure 6). Diffusion-based models showed a steady, linear decline in predictive performance across deciles, with each decile performing slightly worse than the previous one. This differs from functional data, which maintained relatively stable performance across most deciles, with a sharp drop-off occurring in the lowest deciles. More broadly, functional connectomes generally achieved higher prediction performances than structural models for each phenotype across deciles. DTI-based results were replicable when using the entire ABCD sample (n=9371, Supplementary Figure S8) and when predicting a phenotype of higher effect size in the HBN dataset (age, n=767, Supplementary Figure S9).

**Figure 6.**
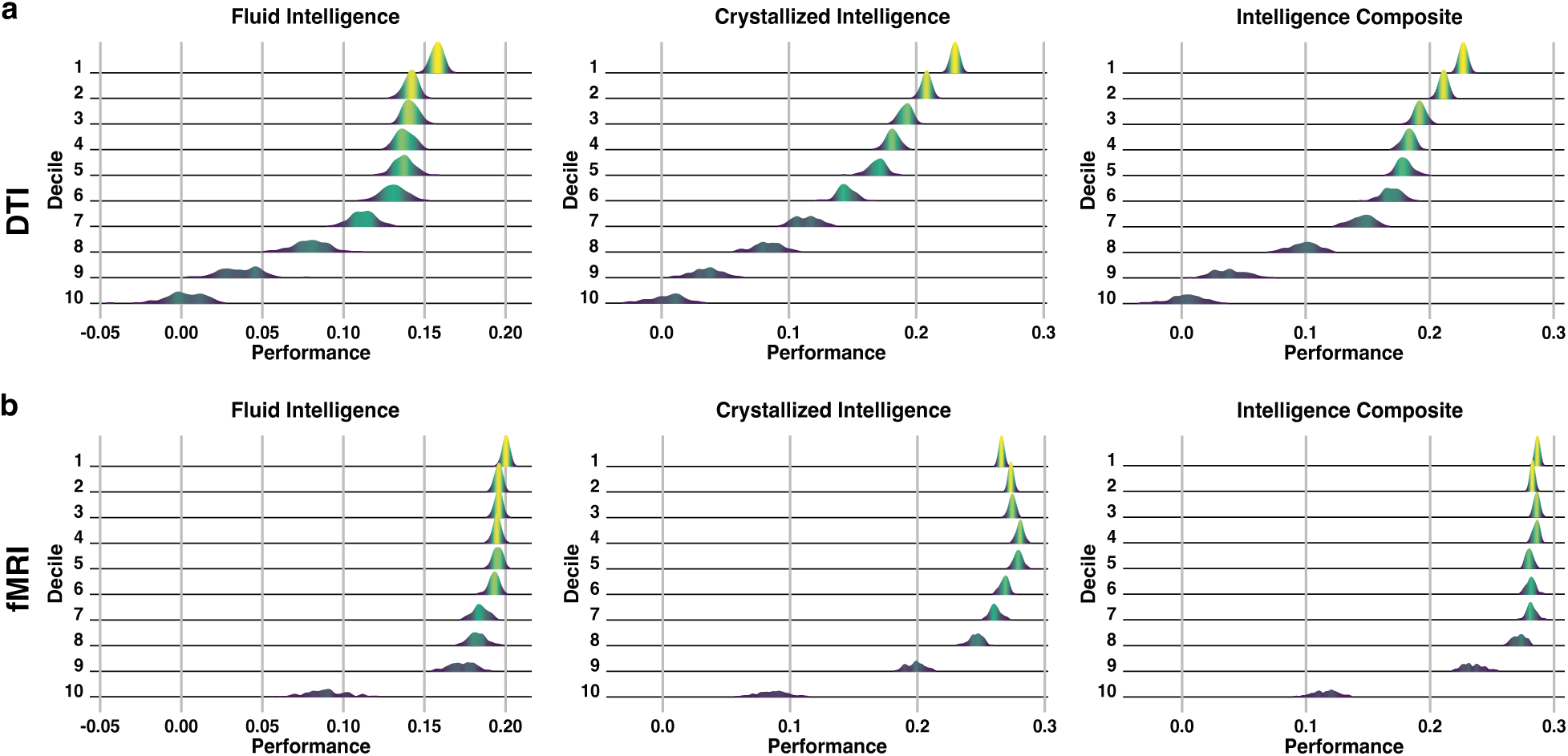
Decile-based predictive modeling performance across DTI and fMRI data in ABCD. Ridgeline plots showing the distribution of prediction performances (Pearson’s r) across 100 iterations for NIH Toolbox age-corrected Fluid, Crystallized, and Composite Intelligence scores using diffusion (A) and fMRI (B) data.

### Sensitivity and Exploratory Analyses

To further evaluate the robustness of our findings, we conducted a series of sensitivity analyses using alternative feature selection and modeling approaches. First, we examined whether the observed predictive performances persisted when using alternative feature groupings. Instead of dividing features into ten deciles, we partitioned them into percentiles (1% subsets), ventiles (5% subsets), and quintiles (20% subsets) based on their correlation with the target phenotype (Supplementary Tables S3, S12, and S13). We also tested whether partitioning features based on alternative statistical metrics influenced results. Features were grouped into non-overlapping bins based on their significance (p-values) or effect size (Pearson’s r-values) with the target phenotype (Supplementary Figure S10). We then evaluated prediction performance while controlling for effects of the non-interest variables age, sex, racial/ethnic minority representation, socioeconomic status, and head motion (Supplementary Table S14). To test the impact of incorporating task fMRI into averaged connectomes on model performances, we repeated PNC analyses using only resting-state connectomes (Supplementary Table 15). In each scenario, features subsets beyond those that are most highly ranked demonstrated significant predictions. Preliminary results also suggest that the top-ranked decile was not always the best-fit model for certain subsets of individuals (Supplementary Figure S11).

## Discussion

Here, we evaluated whether unselected features can meaningfully contribute to building and interpreting predictive models in neuroimaging. Using a decile-based feature ranking paradigm, we demonstrated that significant predictive capabilities are not exclusive to top-ranked connectivity features. Across functional and structural connectivity data and a wide range of phenotypes and datasets, lower-ranked, unselected features achieved significant prediction accuracy often comparable to traditionally selected features comprising the most highly ranked features. Notably, lower-ranked features revealed complementary connectivity patterns that often featured unique neuroanatomical contributions. Overall, the observation that multiple feature sets can achieve significant predictions could lead to divergent neurobiological interpretations of brain-behavior associations. As such, while feature selection may improve many aspects of predictive modeling, it may present only the ‘tip of the iceberg’ when certain disregarded features may be just as meaningful.

Our results are aligned with several emerging neuroimaging trends. First, they support the growing appreciation of widely distributed brain networks, as opposed to more localized or simplified networks. Evidence from human ^39,51–53^ and animal ^54,55^ studies demonstrates that the whole brain responds to stimuli, supporting the view that networks—not single nodes—drive behavior and cognition. Because brain-behavior models depend on widely distributed networks, even edges with weak individual correlations to behavior may play a meaningful role when considered as part of a broader functional network. While sparser models achieved by feature selection are convenient, they may oversimplify the underlying neurobiology of individual variation in behavior.

Next, while predictive models are more replicable than brain-wide association studies ^7,10^, replicability challenges still exist ^5,56,57^. It is not uncommon for multiple of the same phenotype to implicate different neurobiological processes ^19^. Our findings may help explain a portion of this variability. In certain instances of replication failure, the different networks identified across studies may represent select snapshots of a more complex neurobiological landscape rather than true disagreement. This concept aligns with the growing trend toward transparent visualizations, which emphasizes retaining and displaying all data rather than discarding sub-threshold results ^6,58^. Thresholded representations to emphasize differences while obscuring underlying similarities.

Our results also align with recognized differences between multivariate and univariate analyses in neuroimaging ^1,59^. For example, univariate features in the connectome (i.e., edges) tend to have poor reliability ^60,61^. However, multivariate reliability methods (e.g., fingerprinting, discriminability) show comparably high reliability of connectomes ^62–67^. This increased reliability reflects the high-dimensional structure of connectomes. Similarly, selecting a single feature at a time without considering the other features ignores this high-dimensional variance structure ^68^. Such approaches are emerging but remain uncommon^69,70^. Alternatively, approaches that create and select the predictive features, like deep learning, may be paths forward ^71,72^. Overall, the limitations of univariate feature selection should be kept in mind when considering the strengths of predictive modeling.

On the surface, it may appear contradictory that higher and lower-ranked features can predict outcomes similarly. This result is likely explained by the high degree of spatial and temporal autocorrelation among brain features ^49,50^. When adding new features to a model, accuracy will increase only if the new feature has information that is unique from the existing features. Two features that are highly correlated with a phenotype likely will have a high correlation between them. Thus, pooling lower-ranked (but uncorrelated) features may perform as well or better than pooling highly ranked (but highly correlated) features. Additionally, pooling features can enhance robustness by suppressing noise, even if their individual univariate associations are weak. In other words, lower signal-to-noise ratios (SNRs) turn relatively strong aggregate brain-behavior associations into weak univariate associations. Alternatively, predictive models can also inadvertently capture information that is not of interest, such as confounds ^18,73^, and they are known to perform more poorly for individuals who defy sample stereotypes^17^. Lower-ranked features may be more susceptible to learning from such information.

As such, the predictions from lower-ranked features may reflect these properties rather than the underlying neurobiology.

Results from our structural models largely aligned with those generated from functional connectomes; however, while functional model performances were relatively stable until the lowest deciles, diffusion-based model performances showed a more steady, linear decline across deciles. Given that functional connectomes achieved higher prediction performances, one possibility is that the utility of lower deciles persists longer for higher-effect-size predictions. This pattern could suggest that high-effect-size phenotypes are distributed more broadly across the brain, or alternatively, that machine learning is more effective at capturing phenotypes with a larger number of contributing features. However, our initial investigation using age prediction with structural connectomes suggests that this explanation does not hold. An alternative explanation could relate to the low signal-to-noise ratio and high temporal autocorrelation of fMRI time series data, which might yield signal redundancies across deciles and account for why performance remains similar across several deciles. Indeed, preliminary results using functional connectomes generated using partial correlation exhibited a similar pattern of performance decay to that observed with diffusion connectomes.

Nevertheless, considerations discussed for functional models likely hold for structural models.

Identifying and defining biologically meaningful clusters (e.g., subtypes) of individuals within a dataset is of great interest within the field ^74–77^. Our results suggest that multiple neurobiologically distinct models may capture a given phenotype, each requiring different models ^17,78^, which has important implications for these ongoing efforts ^79–88^. For example, exploratory analyses indicate that one subtype may be best predicted by decile 1, while another subtype may be best predicted by decile 2. If this is the case, the top-ranked feature set identified by a particular analysis could be influenced by the relative proportions of subtypes within the sample. As the distribution of subtypes shifts, the features deemed most predictive could also change. While our preliminary results point to this possibility, further analysis is needed to validate the extent to which subtype composition affects which features are selected. Future work could leverage more formal clustering approaches to identify latent subgroups and evaluate whether these align with distinct feature subsets ^89^.

Similarly, unselected, lower-ranked features may be new targets for emerging interventions and strategies for individualized medicine. Traditionally, the highest ranked features from a predictive model are selected as potential targets ^32^, though these features may not match a specific therapy. For example, it is difficult to stimulate subcortical regions with transcranial magnetic stimulation (TMS) ^90,91^. If a model consists of these regions, it would be a poor fit to inform TMS targets. However, overlooked, lower-ranked features may be more anatomically accessible (e.g., cortical versus subcortical) and still predict the phenotype of interest with the same accuracy as the high-ranked features. Looking beyond the traditional way of selecting features may better align predictive models with therapeutic approaches. Still, it remains unclear which features are the best targets for intervention. Brain features that predict inter-individual variation in symptoms or treatment outcomes are not necessarily the same as those causally involved in symptom manifestation ^92,93^.

A strength of this study lies in the comprehensive validation of our models across diverse datasets and methodological frameworks. We leveraged four large-scale neuroimaging datasets to enhance statistical power and generalizability. The PNC, HBN, HCPD, and ABCD datasets exhibit notable variability in key aspects of study design including recruitment geography, behavioral assessment, imaging acquisition, participant demographics, and clinical symptom burden ^25^. Demonstrating the resilience of our models to differences in participant demographics and study design, the utility of commonly disregarded features survived external validation. Neuroimaging predictive models will achieve real-world utility only if they can overcome these dataset-specific idiosyncrasies (i.e., “dataset shifts”) ^25,56,94–96^. Finally, we also performed an extensive set of sensitivity analyses. Our results were robust to both functional and structural imaging modalities and across a range of cognitive, psychiatric, and developmental phenotypic domains.

We used Pearson’s correlation as our main measure of prediction performance. However, many models exhibited negative q^2^, indicating that the sample mean achieves a lower mean square error. Based on q^2^, these models did not successfully predict outcomes. Nevertheless, modes with negative q^2^ can still have high utility. For example, such models may accurately preserve the relative ranking of individuals, which can be valuable for clinical tasks involving risk stratification and prioritization of support or intervention. While a single measure of prediction performance can never fully characterize a model’s performance, our interpretations are based on a single measure, Pearson’s correlation. We present multiple measures to provide a more comprehensive view of the model. Though many models may no longer exhibited significant prediction with q^2^, the main results of the paper still hold.

There were several limitations to our study. First, we pursued a network-centric approach focusing on the connectivity strengths (e.g., edges) between 268 brain regions (e.g., nodes) as features. Node-based models, which may capture local brain dynamics or region-specific activity patterns that are obscured in connectome-based approaches, could reveal different results. However, connectome-based models are more frequently employed in the field, in part due to superior predictive abilities ^97^.

Second, it is unclear whether overlooked features maintain their predictive utility across phenotypes that are hypothesized to rely on more circumscribed functional networks, such as reaction time. Finally, this investigation relies on developmental datasets exclusively from the United States. Future work should investigate the utility of commonly overlooked features in adult populations and those from non-Western countries ^98,99^.

Collectively, our findings highlight that commonly overlooked features may possess comparable predictive and neurobiological significance. They support the growing appreciation of widely distributed brain networks, as opposed to more localized or simplified networks. Our findings also point to the potential presence of subtypes in the connectivity data, wherein different sets of features represent the best model for different groups of individuals. Overall, better understanding and characterization of overlooked features will help improve the generalizability of predictive models.

## Methods

### Datasets

PNC participants were 1291 individuals ages 8-21 recruited from the greater Philadelphia, Pennsylvania area ^43^. All imaging was performed on a single scanner at the Hospital of the University of Pennsylvania. Participants completed rest, emotion task, and n-back task fMRI runs ^100^. Measures of language abilities were the Penn Verbal Reasoning Task from the Penn Computerized Neurocognitive Battery (CNB) and the total standard score from the Wide Range Assessment Test (WRAT) Reading Subscale ^101,102^. Executive function measures were the Letter N-Back, Conditional Exclusion, and Continuous Performance tasks from the CNB.

HBN participants were 1110 individuals ages 6-17 recruited from the New York City, New York region ^40^. Imaging for this multi-site study was performed via the HBN mobile MRI scanner in Staten Island and at the Rutgers University Brain Imaging Center, the CitiGroup Cornell Brain Imaging Center, and the CUNY Advanced Science Research Center. Participants completed two rest fMRI runs as well as ‘Despicable Me’ and ‘The Present’ movie-watching scan sessions. Measures of language abilities were the Elision, Blending Words, Nonword Repetition, Rapid Digit Naming, and Rapid Letter Naming scaled scores from the Comprehensive Test of Phonological Processing (CTOPP-2) and the Phonemic Decoding Efficiency, Sight Word Efficiency, and Total Word Reading Efficiency scaled scores from the Test of Word Reading Efficiency (TOWRE-2) ^103,104^. Executive function measures were the Flanker Inhibitory Control and Attention, List Sorting Working Memory, Pattern Comparison Processing Speed, and Dimensional Change Card Sort age-corrected standard scores from the NIH Toolbox105.

To select developmental and psychiatric phenotypes of meaningful effect size, we first performed predictions using the original CPM framework. We identified scales and subscales with prediction performance *r*>0.10 by testing predictions on the 575 HBN participants who had complete data for all 71 measures. This resulted in 13 scales and subscales. We repeated the predictions using the 747 participants who had data available for all 13 scales. Of these, 6 scales achieved prediction performances of *r*>=0.15, which were used for our decile-based prediction paradigm: the Social Communication Questionnaire (SCQ) ^46^ score, the Hyperactivity Scale score from the Strengths and Difficulties Questionnaire (SDQ) ^106,107^, the Separation Anxiety score from the Screen for Anxiety Related Disorders (SCARED) ^108^, the Learning Problems score from the Conners 3 Self-Report (C3SR) ^109^, and the Internet Addiction Test (IAT) and Parent-Child Internet Addiction Test (PCIAT) ^110^ scores.

HCPD participants were 428 individuals ages 8-22 recruited from St. Louis, Missouri, Twin Cities, Minnesota, Boston, Massachusetts, and Los Angeles, California^42^. Imaging for this multi-site study was performed at Harvard University, the University of California-Los Angeles, the University of Minnesota, and Washington University in St. Louis. Participants completed rest fMRI runs ^111^. Measures of language abilities were the Picture Vocabulary and Oral Reading Recognition age-corrected standard scores from the NIH Toolbox. Executive function measures were the Flanker Inhibitory Control and Attention, List Sorting Working Memory, Pattern Comparison Processing Speed, Dimensional Change Card Sort, and Picture Sequence Memory age-corrected standard scores from the NIH Toolbox.

ABCD participants were 9371 individuals ages 9-10 recruited from 21 sites across the United States: Children’s Hospital Los Angeles, Florida International University, Laureate Institute for Brain Research, Medical University of South Carolina, Oregon Health & Science University, SRI International, UC San Diego, UCLA, University of Colorado Boulder, University of Florida, University of Maryland at Baltimore, University of Michigan, University of Minnesota, University of Pittsburgh, University of Rochester, University of Utah, University of Vermont, University of Wisconsin-Milwaukee, Virginia Commonwealth University, Washington University in St. Louis, and Yale University ^41,112^. Cognitive measures were the Fluid Intelligence, Crystalized Intelligence, and Intelligence Composite age-corrected scores from the NIH Toolbox.

### fMRI data preprocessing

In all datasets, data were motion-corrected. Additional preprocessing steps were performed using BioImage Suite ^113^. This included regression of covariates of no interest from the functional data, including linear and quadratic drifts, mean cerebrospinal fluid signal, mean white matter signal, and mean global signal. Additional motion control was applied by regressing a 24-parameter motion model, which included six rigid body motion parameters, six temporal derivatives, and the square of these terms, from the data. Subsequently, we applied temporal smoothing with a Gaussian filter (approximate cutoff frequency=0.12 Hz) and gray matter masking, as defined in common space. Then, the Shen 268-node atlas was applied to parcellate the denoised data into 268 nodes ^114^. Finally, we generated functional connectivity matrices by correlating each node time series data pair and applying the Fisher transform.

Data were excluded for poor data quality, missing nodes due to lack of full brain coverage, high motion (>0.2 mm mean frame-wise motion), or missing behavioral/phenotypic data. Each participant’s connectome included all available resting-state and task fMRI data with low motion (<0.2 mm). Connectomes for individual conditions (i.e., resting-state and task) were created independently and then averaged. Combining connectomes across fMRI data improves reliability and predictive power ^15,115^. Participants without one low-motion fMRI run were excluded. For PNC, 246 participants were excluded due to image quality or motion, and 61 participants were excluded due to incomplete phenotypic data. For HBN, 1387 participants were excluded due to image quality or motion, and 829 participants were excluded due to incomplete phenotypic data. For HCPD, 57 participants were excluded due to image quality or motion, and 167 participants were excluded due to incomplete phenotypic data. For ABCD, 342 participants were excluded due to incomplete phenotypic data.

### ABCD diffusion data preprocessing

FIB files were downloaded from https://brain.labsolver.org. As described on https://brain.labsolver.org/, a multishell diffusion scheme was used, and the b-values were 500, 1000, 2000, and 3000 s/mm². The number of diffusion sampling directions were 6, 15, 15, and 60, respectively. The in-plane resolution was 1.7 mm. The slice thickness was 1.7 mm. The diffusion MRI data were rotated to align with the AC-PC line at an isotropic resolution of 1.7 (mm). The restricted diffusion was quantified using restricted diffusion imaging ^116^. The diffusion data were reconstructed using generalized q-sampling imaging ^117^ with a diffusion sampling length ratio of 1.25. The tensor metrics were calculated using DWI with b-value lower than 1750 s/mm².

As stated in previous works and duplicated here for consistency ^118,119^, whole-brain fibre tracking was conducted with DSI-Studio with quantitative anisotropy (QA) as the termination threshold. QA values were computed in each voxel in their native space for each subject and were then used to warp the brain to the template in Montreal Neurological Institute (MNI) space using the statistical parametric mapping nonlinear registration algorithm. Once in MNI space, spin density functions were again reconstructed with a mean diffusion distance of 1.25 mm using three fibre orientations per voxel. Fibre tracking was performed in DSI-Studio with an angular cut-off of 60 degrees, step size of 1.0 mm, minimum length of 30 mm, spin density function smoothing of 0.0, maximum length of 300 mm and a QA threshold determined by the diffusion-weighted imaging signal. Deterministic fibre tracking using a modified FACT algorithm ^120^ was performed until 10,000,000 streamlines were reconstructed for each individual. We used the Shen atlas^114^ in MNI space with 268 nodes to construct individual structural connectomes: the pairwise connectivity strength was calculated as the average QA value of each fibre connecting the two end regions and thresholded at 0.001, which results in a 268 × 268 adjacency matrix for each participant.

### Creating latent factors of language abilities and executive function

Whether machine learning models are used for real-world prediction or advancing our understanding of neurobiology ^121,122^, it is important to overcome dataset-specific idiosyncrasies ^1^. The PNC, HCPD, and HBN datasets employed disparate measures to assess executive function and language abilities. To harmonize these measures and facilitate direct comparisons across datasets, we used principal component analysis (PCA) to derive’latent’ factors of executive function and language abilities within each dataset. Briefly, as in previous work ^25^, PCA was applied to behavioral measures to reduce measurement noise and derive a single composite factor per domain. For PNC and HBN, PCA was conducted using participants without usable neuroimaging data to maintain independent train and test groups, while for HCPD, PCA was performed within each fold of cross-validation, using only training data. Behavioral data from participants with imaging data were projected onto the first principal component, which explained 70%, 55%, and 77% of the variance in language measures and 53%, 48%, and 40% in EF measures for PNC, HBN, and HCPD, respectively.

### Decile-based connectome-based predictive modeling

We implemented an original adaptation of the original CPM framework^44^ to evaluate the predictive utility of connectivity features across varying levels of association with the target phenotype. For within-dataset predictions, we performed 100 iterations of a 10-fold cross validation scheme. Within each fold, the model was trained in approximately 90% of the data and tested in the remaining 10% of the data. For each target phenotype, within each fold, we computed the Pearson correlation coefficient between every connectivity edge and the target phenotype across all participants in the training subset. This univariate analysis provided a measure of association strength for each feature. Features were then ranked in descending order based on the absolute value of their correlation coefficients, ensuring that both positive and negative associations were considered equally in the ranking process. Following ranking, the complete set of connectivity features was partitioned into ten non-overlapping deciles. Each decile represented 10% of the total features, with decile 1 comprising the top 10% of features exhibiting the strongest associations with the target phenotype, and decile 10 containing the bottom 10%. Predictions were performed using each decile and tested on the fold’s test set. For sensitivity analyses, independent predictive models were created to account for non-interest variables (age, sex, racial/ethnic minority representation, socioeconomic status, head motion, and clinical symptom burden) via partial correlation at the step in which brain edges were related to each phenotype ^123^.

### Ridge regression connectome-based predictive modeling

We repeated analyses using ridge regression CPM to better suit the high-dimensional nature of connectivity data ^15^. Specifically, due to the positive semi-definite nature of a functional connectivity matrix, the edges are not independent. Ridge regression is more robust than OLS in this case. Instead of summing selected edges and fitting a one-dimensional OLS model, we directly fit a ridge regression model with training individuals using the selected edges from all the tasks and apply the model to testing individuals in the cross-validation framework. The L2 regularization λ parameter was chosen using Bayesian optimization over the training data within each fold.

Specifically, MATLAB’s ‘fitrlinear’ function was used with automatic hyperparameter optimization to select the λ value that minimized the cross-validated mean squared error on the training set. All hyperparameter tuning was performed strictly within the training data to avoid information leakage.

### Model performance

Predictive models were trained and tested within each dataset using 100 iterations of 10-fold cross-validation. Model performance was evaluated with Pearson’s correlation (r), representing the correspondence between predicted and actual behavioral scores. We report mean Pearson’s r across the 100 iterations, along with the cross-validation coefficient of determination (q2) and mean square error (MSE) ^5^.

Within-dataset plots show data for the median-performing models. To generate null distributions for significance testing, we randomly shuffled the correspondence between behavioral variables and connectivity matrices 1000 times and re-ran the CPM analysis with the shuffled data. Based on these null distributions, the p-values for predictions were calculated as in prior work ^124^. Only a positive association between predicted and actual values indicates prediction above chance (with negative associations indicating a failure to predict), so one-tailed p-values are reported. Pearson’s correlation was tested between actual and predicted values to evaluate cross-dataset predictions.

### Model contribution

Predictive networks identified using CPM or ridge regression are composed of multiple brain regions and networks. Each node is defined as a single region in the 268-node Shen atlas, and each node is associated with multiple features (specifically, the edges connecting it to the other 267 nodes). To quantify the contribution of each node and network, we first identified the median-performing model across the 100 iterations for each phenotype. All interpretability analyses were based on this median model.

For ridge regression models, we used the mean of each edge’s coefficient across the 10 folds of the median-performing model. The Haufe transform^48^ was applied to model weights. Node contributions were calculated by summing the absolute value of Haufe-transformed weights across all edges connected to a given node. Network-level contributions were computed by averaging the absolute value of Haufe-transformed weights within or between canonical functional networks, normalized by network size.

Specifically, to quantify the contribution of each node to a given predictive model, we calculated the 𝑛^th^ node’s weight summed across all edges (labeled 𝑊_n_) to the model as: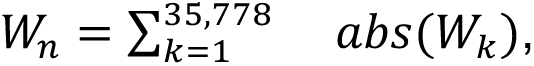 for all 𝑘 edges connected to the 𝑛^th^ node. Next, for the network level, 𝑊_k_ was averaged over each edge within or between canonical functional networks.

For CPM models, edge selection is inherently binary. Each edge is either selected (1) or not (0) in a given fold. We averaged these binary masks across the 10 folds of the median-performing model. Node and network contributions were calculated as with ridge, using these fold-averaged values.

### Decile feature comparisons

To quantify the shared variance of model weights across feature deciles, we computed pairwise squared Pearson correlations (r²) between decile models. For node-level comparisons, ridge regression edge weights were summed for each of the 268 regions defined by the Shen atlas, resulting in a 268-dimensional vector of node-wise importance values per model. The r² values between these vectors indicate the proportion of variance in node-level weight distributions shared between models across deciles. For network-level comparisons, edge-wise weights were averaged within each of 55 canonical functional networks, producing a network-level weight vector for each model. Pairwise r² values were then calculated between these vectors across deciles, reflecting the degree of similarity in the network-level distribution of feature weights.

For connectome-based predictive modeling (CPM), a similar approach was used, except that binary selection masks (1 = selected, 0 = not selected) were used in place of continuous weights. Node-wise importance vectors were created by summing binary edge selections per node, and network-level vectors were generated by averaging binary selections within each network. Squared correlations (r²) between these binarized vectors were then calculated across deciles to estimate the similarity of features.

## Data availability

Data are available through the Philadelphia Neurodevelopmental Cohort, Human Connectome Project in Development, Healthy Brain Network, and Adolescent Brain Cognitive Development Study datasets.

## Code availability

Analyses were conducted using Matlab R2024a. Code is available via GitHub at https://github.com/brendan-adkinson/overlooked-features. Preprocessing was carried out using Bioimage Suite v.3.01, which is freely available (https://medicine.yale.edu/bioimaging/suite/). Additional preprocessing was performed with the Human Connectome Project minimal preprocessing pipeline v.3.4.0 (https://github.com/Washington-University/HCPpipelines/releases).

## Acknowledgements

This study was supported by the National Institute on Minority Health and Health Disparities under grant 1F30MD018941 (obtained by B.D.A.). B.D.A. was also supported by NIH Medical Scientist Training Program Training Grant T32GM136651.

M.R. was supported by the National Science Foundation Graduate Research Fellowship under grant DGE2139841. L.T. was supported by the Gruber Science Fellowship. S.N. was supported by the National Institute of Mental Health under grant K00MH122372. The funders had no role in study design, data collection and analysis, decision to publish, or preparation of the manuscript. Any opinions, findings, and conclusions or recommendations expressed in this material are those of the authors and do not necessarily reflect those of the funding agencies. The authors thank Rubi Ching for her insightful feedback and assistance with figure design. They also thank Karis Gillen and Lucy Yang for their assistance with table preparation.

## Author contributions statement

**Brendan D. Adkinson:** Writing – review & editing, Writing – original draft, Visualization, Resources, Methodology, Investigation, Formal analysis, Data curation, Funding acquisition, Conceptualization. **Matthew Rosenblatt:** Writing – review & editing, Resources, Methodology, Investigation, Formal analysis. **Huili Sun**: Resources. **Javid Dadashkarimi:** Resources, Methodology, Data curation. **Link Tejavibulya:** Data curation. **Corey Horien**: Writing – review & editing. **Margaret L. Westwater**: Writing – review & editing. **Raimundo X. Rodriguez**: Resources, Formal analysis. **Stephanie Noble:** Writing – review & editing, Methodology, Conceptualization. **Dustin Scheinost:** Writing – review & editing, Writing – original draft, Supervision, Software, Resources, Project administration, Methodology, Investigation, Funding acquisition, Conceptualization.

