## Supplement for "Overlooked features lead to divergent neurobiological interpretations of brain-based machine learning biomarkers"

**Table of contents**

Figure S1. PNC null distributions
Figure S2. HCPD null distributions
Figure S3. HBN null distributions
Figure S4. Network- and node-level features across deciles for PNC, HCPD, and HBN
Figure S5. Variance explained between CPM decile model features at the network and node levels across PNC, HCPD, and HBN
Figure S6. Network similarity across deciles using edges present in at least five of the ten cross validation folds
Figure S7. Variance explained between ridge decile and full-connectome model features at the network and node levels across PNC, HCPD, and HBN
Figure S8. Decile-based predictive modeling performance for DTI across the entire ABCD sample
Figure S9. HBN predictions of age using DTI and fMRI across deciles
Figure S10. Prediction performances across binned r and p values
Figure S11. Deciles feature subsets in which participants' 'best fit' model was achieved
Table S1. Characteristics of the PNC, HBN, HCPD, and ABCD datasets (adapted from Adkinson et al., 2024)
Table S2. Paired t-tests comparing decile 1 to other deciles across executive function and language models
Table S3. Within-dataset cognitive phenotype predictions across 1%, 5%, 10%, and 20% of features
Table S4. Cross-dataset performances testing in PNC
Table S5. Cross-dataset performances testing in HCPD
Table S6. Cross-dataset performances testing in HBN
Table S7. Edge selection across folds for each decile-based model
Table S8. Ridge regression cognitive phenotype predictions using all features and decile-based subsets
Table S9. Paired t-tests for ridge regression comparing decile 1 to other deciles across executive function and language models
Table S10. Partial correlation CPM predictions across 1%, 5%, 10%, and 20% of features
Table S11. Partial correlation ridge regression predictions
Table S12. Psychiatric and developmental predictions across 1%, 5%, 10%, and 20% of features
Table S13. Age and sex predictions across 1%, 5%, 10%, and 20% of features
Table S14. Prediction performances controlling for confounds
Table S15. PNC rest connectome predictions

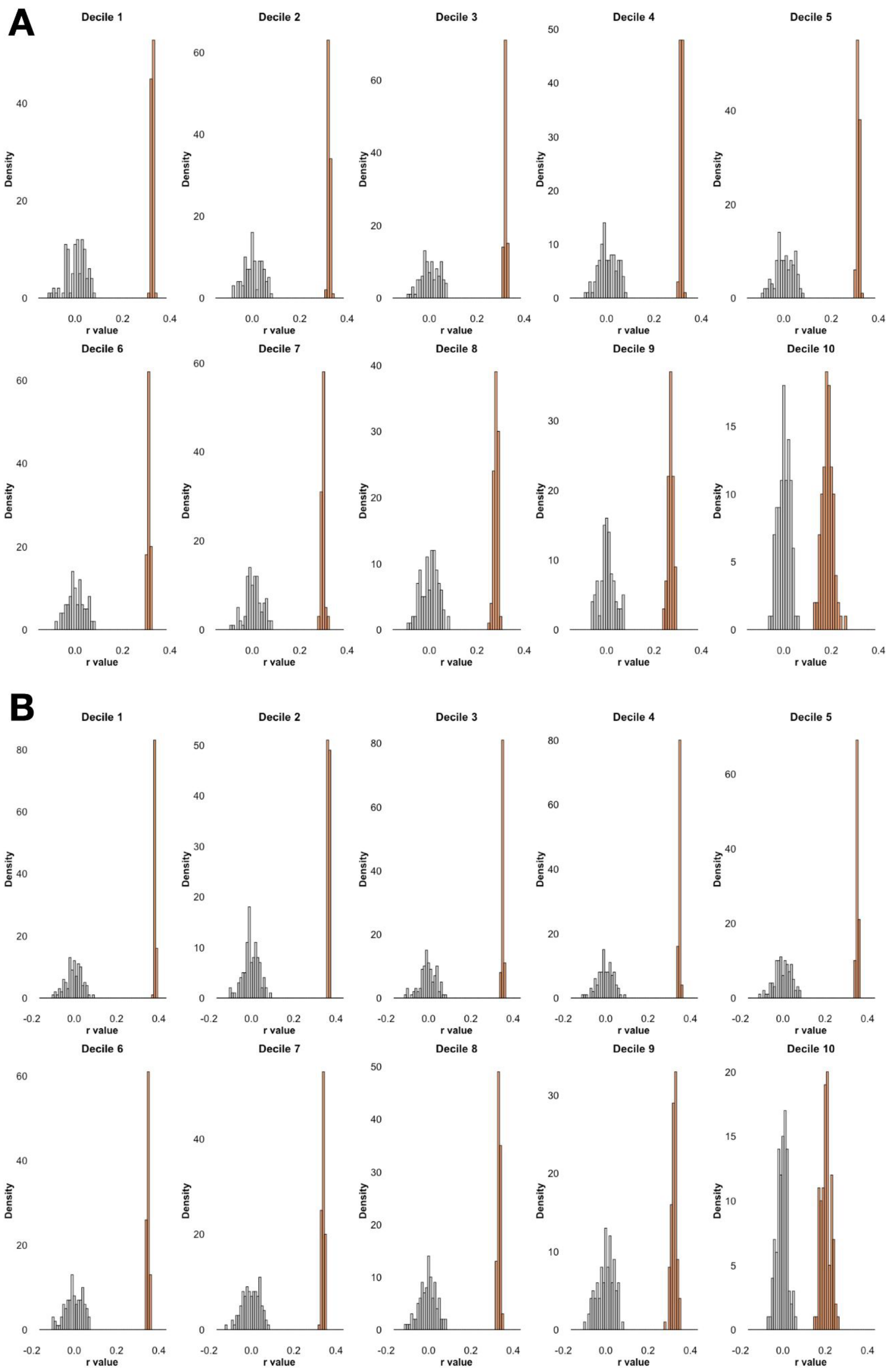


**Figure S1. PNC null distributions.** 100 iterations of real data (orange) next to 1000 iterations of null data (gray) for executive function (A) and language abilities (B).

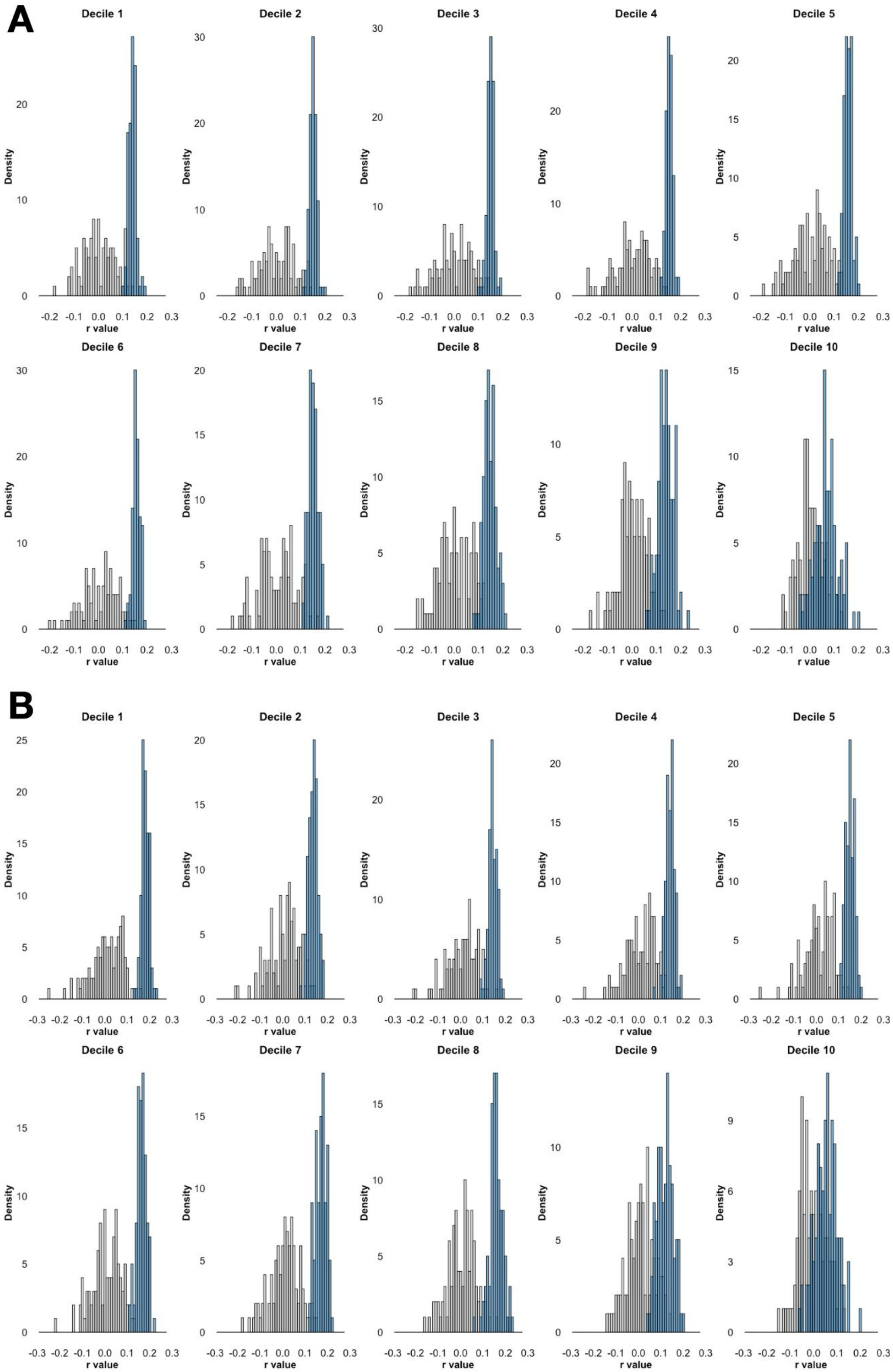


**Figure S2. HCPD null distributions.** 100 iterations of real data (blue) next to 1000 iterations of null data (gray) for executive function (A) and language abilities (B).

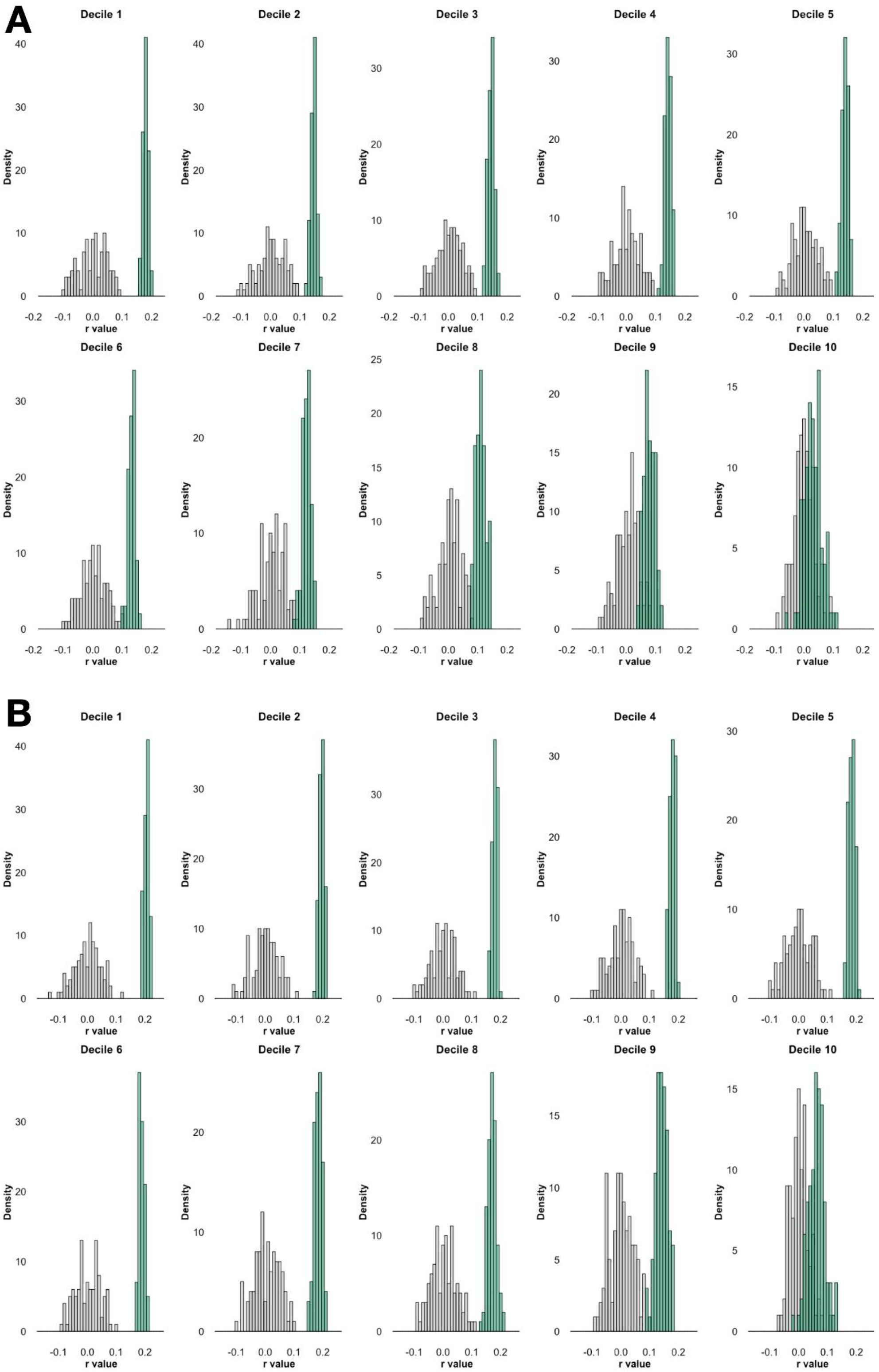


**Figure S3. HBN null distributions.** 100 iterations of real data (green) next to 1000 iterations of null data (gray) for executive function (A) and language abilities (B).

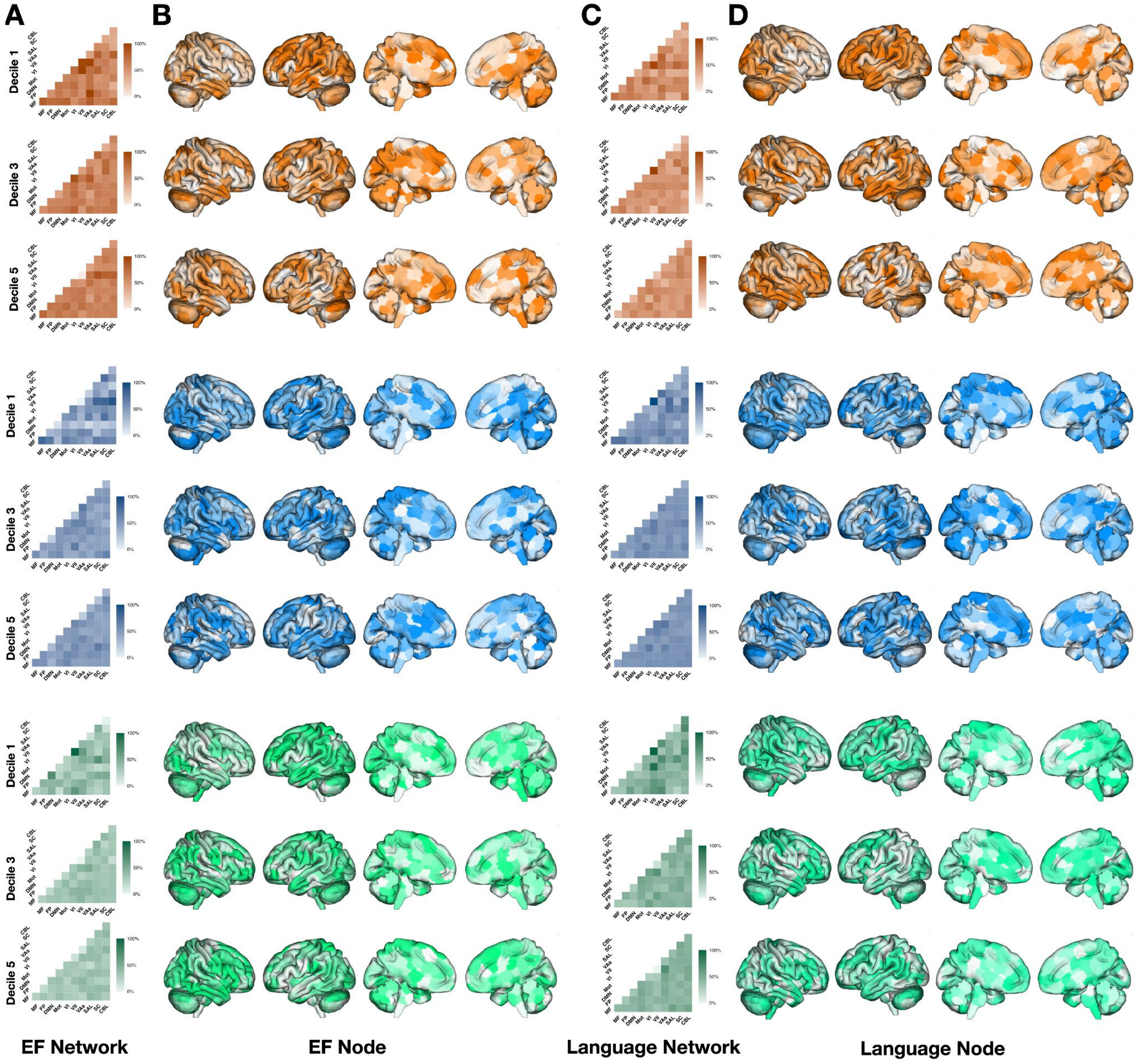


**Figure S4. Network- and node-level features across deciles for PNC, HCPD, and HBN.** Canonical network contributions to predicted executive function (A) and language abilities (C) across PNC (orange), HCPD (blue), and HBN (green). Contributions represent the number of selected features averaged across folds (i.e., proportion of folds in which each edge was selected in the median-performing model) grouped by canonical functional network pairs. Diagonal cells represent contributions of edges within a single network; off-diagonal cells represent contributions of edges between networks. Values were normalized by the number of edges in each network group. Darker colors indicate higher relative contribution. Network Labels: MF, medial frontal; FP, frontoparietal; DMN, default mode; Mot, motor cortex; VI, visual A; VII, visual B; VAs, visual association; SAL, salience; SC, subcortical; CBL, cerebellum. Node-level contributions to executive function (B) and language abilities (D) for PNC (orange), HCPD (blue), and HBN (green) predictions.

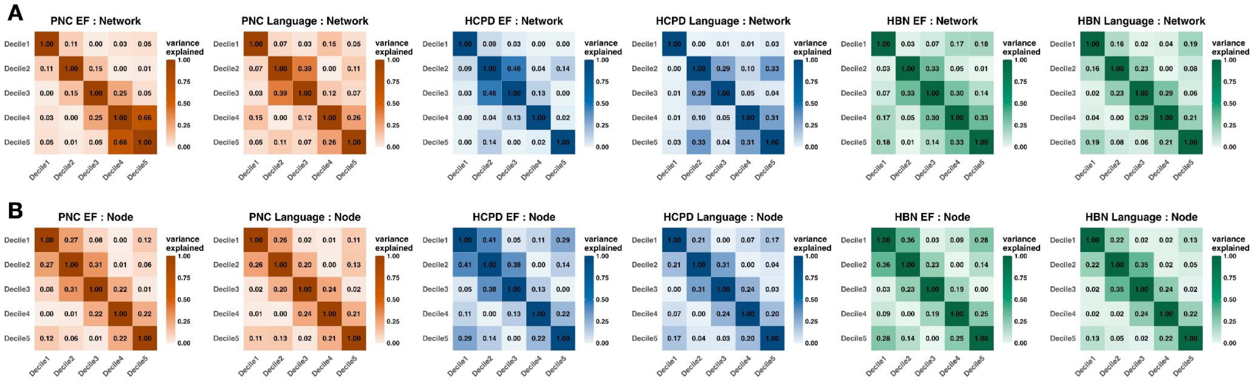


**Figure S5. Variance explained between CPM decile model features at the network and node levels across PNC (orange), HCPD (blue), and HBN (green).** (A) Pairwise variance explained between node degree vectors across deciles. (B) Pairwise variance explained between network-level feature distributions across deciles. Higher variance explained values reflect greater similarity in features across deciles.

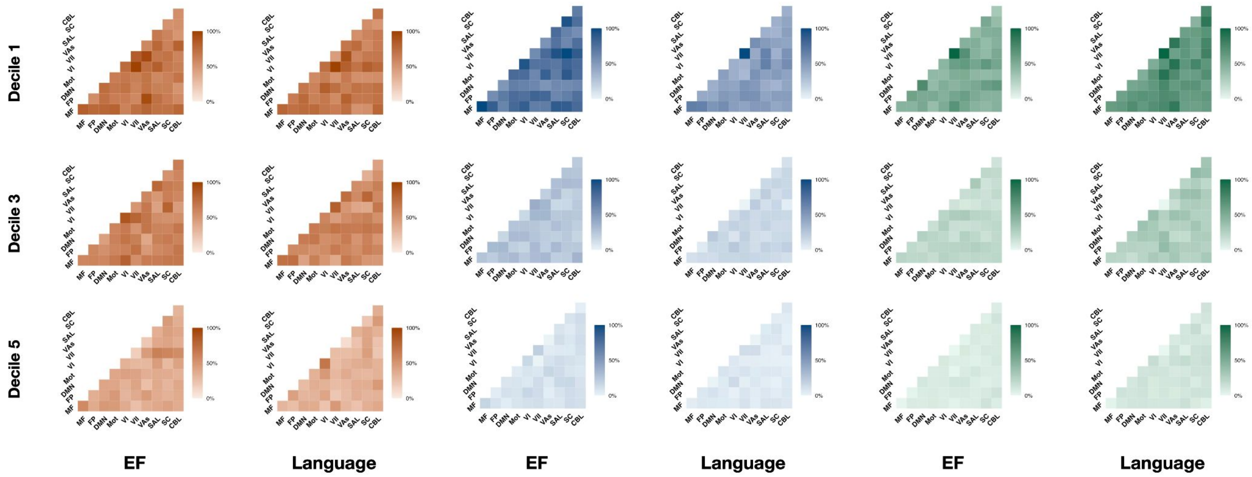


**Figure S6. Network similarity across deciles using edges present in at least five of the ten cross validation folds.** As an alternative to main text figures which utilize the mean of folds, network patterns are shown for PNC (orange), HCPD (blue), and HBN (green).

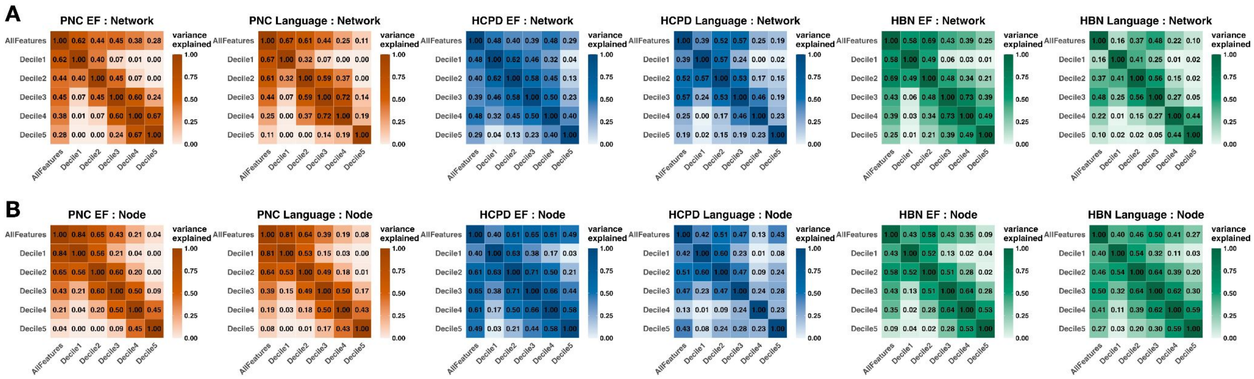


**Figure S7. Variance explained between ridge decile and full-connectome model features at the network and node levels across PNC (orange), HCPD (blue), and HBN (green).** (A) Pairwise variance explained ($r^2$) between node-level weight vectors across deciles and the all-features model. Each value represents the squared Pearson correlation between node-wise sums of ridge regression weights for two models, calculated across 268 regions in the Shen atlas. For each model, edge-wise weights were summed per node, generating a 268-dimensional vector. (B) Pairwise variance explained ($r^2$) between network-level weight distributions across deciles and the all-features model. For each ridge regression model, edge-wise weights were averaged within 55 canonical functional networks, and the resulting network-level vectors were compared using squared Pearson correlation. Higher $r^2$ values indicate greater similarity in node- or network-level feature importance between models.

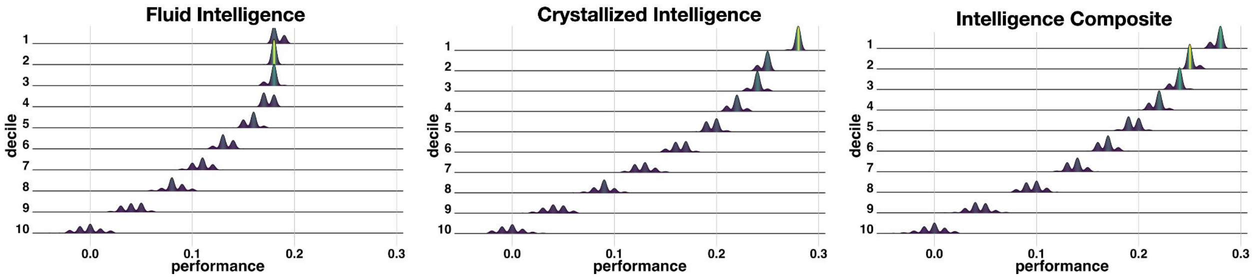


**Figure S8. Decile-based predictive modeling performance for DTI across the entire ABCD sample (n=9371).** Ridgeline plots showing the distribution of prediction performances (Pearson's r) across 100 iterations for NIH Toolbox age-corrected Fluid, Crystallized, and Composite Intelligence scores.

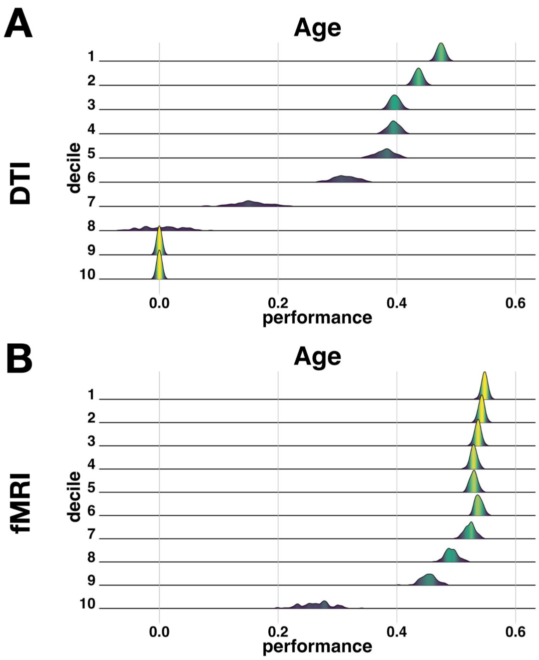


**Figure S9. HBN predictions of age using DTI (A) and fMRI (B) across deciles.** Ridgeline plots show the distribution of prediction performances (Pearson's r) across 100 iterations.

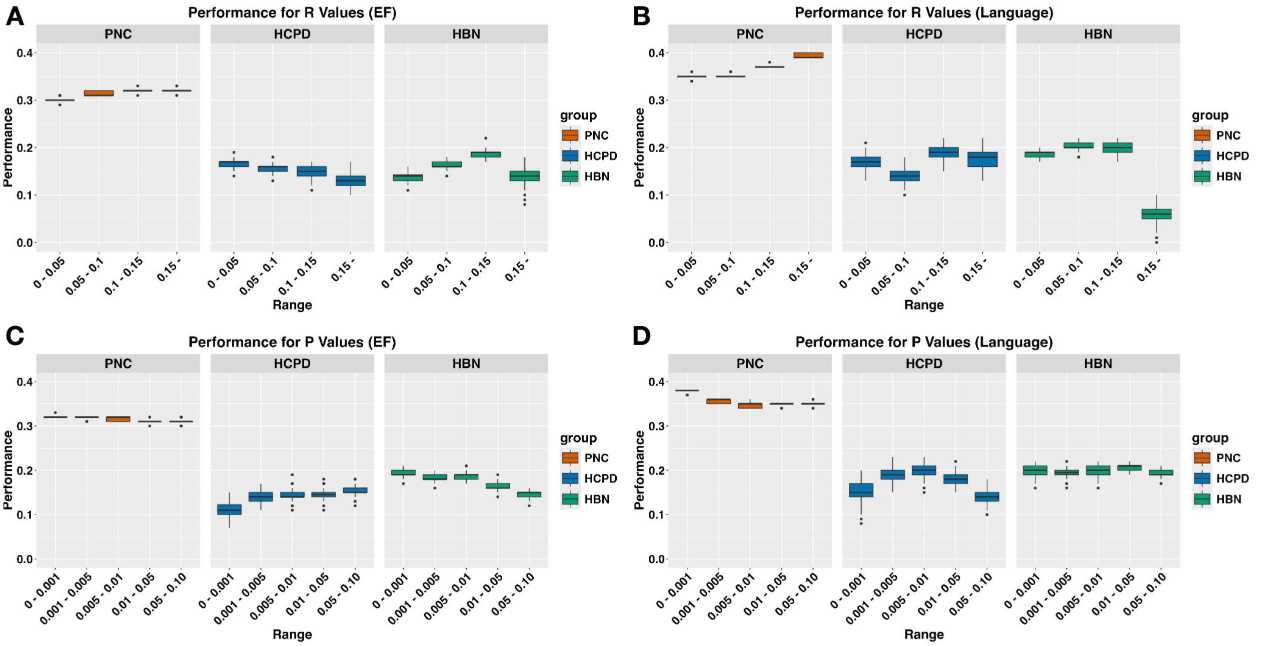


**Figure S10. Prediction performances across binned r and p values.** We evaluated model performance using subsets of features grouped by the significance and effect size of their univariate correlation with executive function (EF). The purpose of this analysis was to evaluate how common feature selection thresholds influence predictive modeling. (A) Features were binned based on their Pearson r-values (e.g., 0–0.05, 0.05–0.10, etc.) and used to predict EF in PNC (orange), HCPD (blue), and HBN (green). (C) Features were binned based on their p-values (e.g., common thresholds of p<0.001, 0.001–0.005, etc.) and used similarly. This binning approach isolates feature subsets of varying statistical strength without relying on rank-based deciles. Each model was trained using only the features within the specified bin, and results reflect average cross-validated performance across 100 iterations.

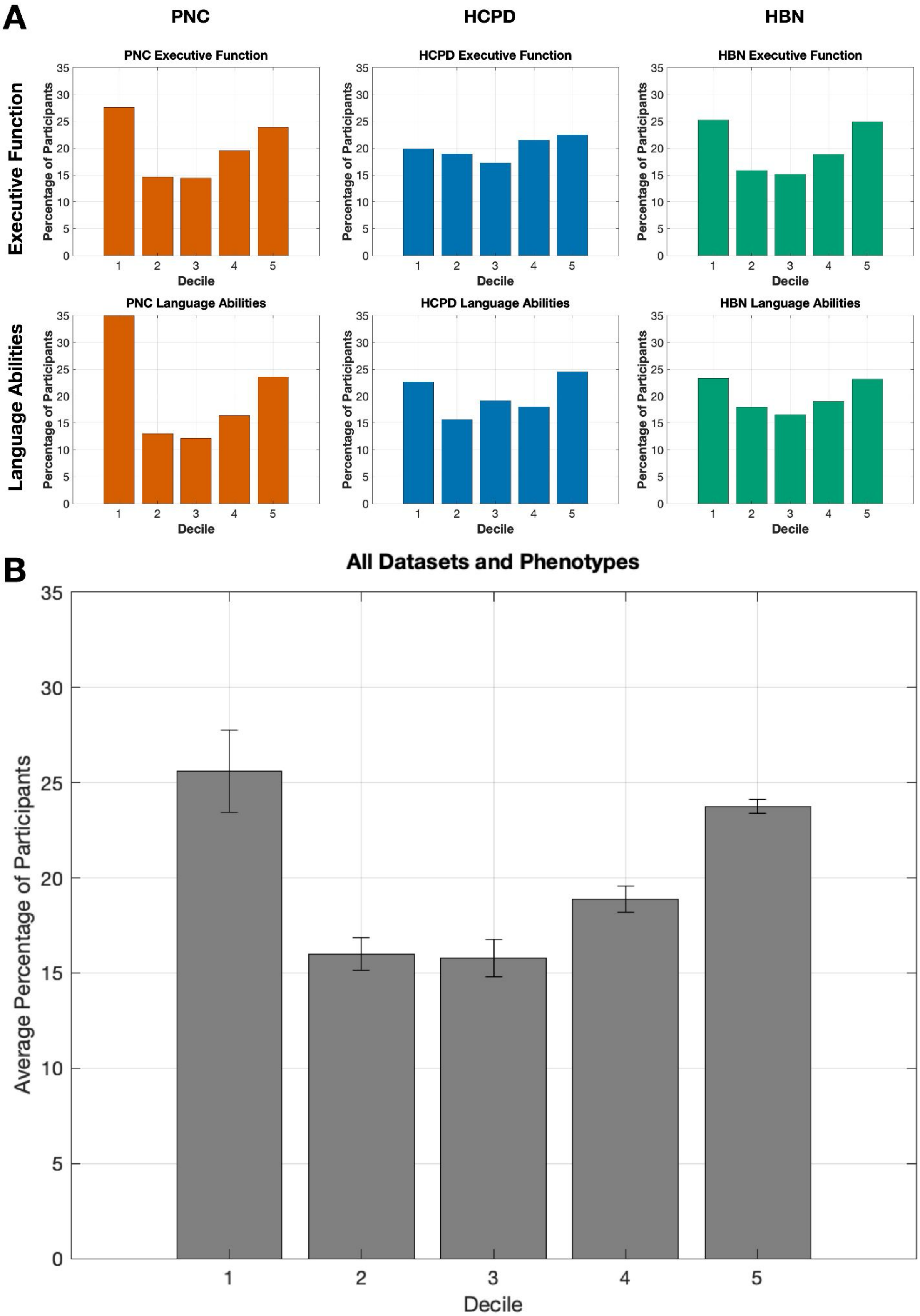


**Figure S11. Deciles feature subsets in which participants' 'best fit' model was achieved.** Our results suggest that a given phenotype may have multiple neurobiologically distinct predictive models, possibly reflecting the existence of subtypes, or groups of individuals who differ in underlying neurobiology and therefore require different models for accurate prediction. We explored this possibility by investigating which decile feature subsets produced the lowest prediction error for participants (i.e., their ‘best fit’ model). (A) Bar charts show the percentage of participants whose absolute prediction error was smallest in each decile. Error was defined at the individual participant level as the absolute discrepancy between a model’s predicted and observed value. A higher percentage in a given decile suggests that more participants had their most accurate predictions (lowest error) in that decile relative to the other deciles. (B) Averaged across PNC executive function, PNC language, HCPD executive function, HCPD language, HBN executive function, and HBN language. Plot error bars show standard error of the mean.

| Dataset | Covariate | Mean | Median | IQR | Percent |
|---|---|---|---|---|---|
| PNC | Age | 14.58 years | 15.00 years | 5.00 years | |
| | Sex (male) | | | | 46.79% |
| | Racial/Ethnic Minority Representation | | | | 52.52% |
| | Socioeconomic Status | N/A | N/A | N/A | |
| | Head Motion | 0.07 mm | 0.06 mm | 0.05 mm | |
| | Clinical Symptom Burden | | | | 73.90% |
| | Imaging Study Design | Single-Site (Hospital of the University of Pennsylvania) | | | |
| | fMRI Tasks | Rest 1, Emotion Task, N-Back Task | | | |
| | Executive Function Tasks | Penn CNB Letter N-Back, Conditional Exclusion, Continuous Performance | | | |
| | Language Abilities Tasks | Penn CNB Verbal Reasoning, Wide Range Assessment Test (WRAT) Reading Subscale | | | |
| HBN | Age | 10.47 years | 10.08 years | 3.91 years | |
| | Sex (male) | | | | 64.05% |
| | Racial/Ethnic Minority Representation | | | | 51.53% |
| | Socioeconomic Status | 50.98 Barratt score | 54.50 Barratt score | 15.50 Barratt score | |
| | Head Motion | 0.11 mm | 0.11 mm | 0.05 mm | |
| | Clinical Symptom Burden | | | | 91.98% |
| | Imaging Study Design | Multi-Site (HBN mobile MRI scanner in Staten Island, Rutgers University Brain Imaging Center, CitiGroup Cornell Brain Imaging Center, CUNY Advanced Science Research Center) | | | |
| | fMRI Tasks | Rest 1, Rest 2, 'Despicable Me' movie-watching, 'The Present' movie-watching | | | |
| | Executive Function Tasks | NIH Toolbox Flanker Inhibitory Control and Attention, List Sorting Working Memory, Pattern Comparison Processing Speed, Dimensional Change Card Sort | | | |
| | Language Abilities Tasks | Comprehensive Test of Phonological Processing (CTOPP-2) Elision, Blending Words, Nonword Repetition, Rapid Digit Naming, Rapid Letter Naming<br>Test of Word Reading Efficiency (TOWRE-2) Phonemic Decoding Efficiency, Sight Word Efficiency, Total Word Reading Efficiency | | | |
| HCPD | Age | 14.78 years | 14.58 years | 6.58 years | |
| | Sex (male) | | | | 47.66% |
| | Racial/Ethnic Minority Representation | | | | 46.03% |
| | Socioeconomic Status | $136,626.45 family income | $125,000.00 family income | $82,000.00 family income | |
| | Head Motion | 0.08 mm | 0.07 mm | 0.04 mm | |
| | Clinical Symptom Burden | | | | N/A |
| | Imaging Study Design | Multi-Site (Harvard University, University of California-Los Angeles, University of Minnesota, Washington University in St. Louis) | | | |
| | fMRI Tasks | Rest 1 | | | |
| | Executive Function Tasks | NIH Toolbox Picture Vocabulary, Oral Reading Recognition | | | |
| | Language Abilities Tasks | NIH Toolbox Flanker, List Sorting Working Memory, Pattern Comparison Processing Speed, Dimensional Change Card Sort, Picture Sequence Memory | | | |
| ABCD (sample with fMRI & DTI) | Age | 9.95 years | 10.00 years | 1.08 years | |
| | Sex (male) | | | | 49.82% |
| | Racial/Ethnic Minority Representation | | | | 41.88% |
| | Socioeconomic Status | <$5K: 2.58%, $5K-$11,999: 3.06%, $12K-$15,999: 1.85%, $16K-$24,999: 3.62%, $25K-$34,999: 5.01%, $35K-$49,999: 7.56%, $50K-$74,999: 13.14%, $75K-$99,999: 14.58%, $100K-$199,999: 30.55%, ≥$200K: 10.56% family income | | | |
| | Head Motion (fMRI) | 0.12 mm | 0.11 mm | 0.05 mm | |
| | Clinical Symptom Burden (CBCL Problems T) | 45.41 | 45.00 | 14.00 | |
| | Imaging Study Design | Multi-Site (Children's Hospital Los Angeles, Florida International University, Laureate Institute for Brain Research, Medical University of South Carolina, Oregon Health & Science University, SRI International, UC San Diego, UCLA, University of Colorado Boulder, University of Florida, University of Maryland at Baltimore, University of Michigan, University of Minnesota, University of Pittsburgh, University of Rochester, University of Utah, University of Vermont, University of Wisconsin-Milwaukee, Virginia Commonwealth University, Washington University in St. Louis, Yale University) | | | |
| | fMRI Tasks | Rest 1, Rest 2, Rest 3, Rest 4 | | | |
| | Cognitive Tasks | NIH Toolbox Fluid Intelligence, Crystalized Intelligence, Intelligence Composite | | | |

**Table S1. Characteristics of the PNC, HBN, HCPD, and ABCD datasets (adapted from Adkinson et al., 2024[25]).**

| Comparison | p value |
|---|---|
| Decile 1 vs. Decile 2 | 0.11 |
| Decile 1 vs. Decile 3 | 0.04 |
| Decile 1 vs. Decile 4 | 0.03 |
| Decile 1 vs. Decile 5 | 0.07 |
| Decile 1 vs. Decile 6 | 0.11 |
| Decile 1 vs. Decile 7 | 0.41 |
| Decile 1 vs. Decile 8 | 0.03 |
| Decile 1 vs. Decile 9 | 0.01 |
| Decile 1 vs. Decile 10 | 0.00 |

**Table S2. Paired t-tests comparing decile 1 to other deciles across executive function and language models.** To assess whether decile 1 features yielded significantly better prediction performance than features from other deciles, we conducted paired t-tests comparing decile 1 to each of deciles 2 through 10. The comparison was performed across the six primary models (executive function and language phenotypes for PNC, HBN, and HCPD). Each p-value reflects a paired t-test comparing the prediction accuracy (e.g., Pearson's r) of decile 1 versus another decile across the six models.

| Phenotype | Percent of Edges | Percentile | Performance | | Percent of Edges | Ventile | Performance | | Percent of Edges | Decile | Performance | | Percent of Edges | Quintile | Performance | |
|---|---|---|---|---|---|---|---|---|---|---|---|---|---|---|---|---|
| | | | r | q2 | | | r | q2 | | | r | q2 | | | r | q2 |
| | | 1 | 0.32 | 0.09 | | 1 | 0.33 | 0.09 | | 1 | 0.33 | 0.09 | | 1 | 0.33 | 0.09 |
| | | 10 | 0.32 | 0.08 | | 2 | 0.32 | 0.08 | | 2 | 0.32 | 0.08 | | 2 | 0.32 | 0.08 |
| | | 20 | 0.32 | 0.08 | | 4 | 0.32 | 0.08 | | 3 | 0.32 | 0.08 | | 3 | 0.31 | 0.08 |
| | | 30 | 0.31 | 0.07 | | 6 | 0.32 | 0.08 | | 4 | 0.31 | 0.08 | | 4 | 0.29 | 0.06 |
| | | 40 | 0.30 | 0.07 | | 8 | 0.31 | 0.08 | | 5 | 0.31 | 0.08 | | 5 | 0.27 | 0.05 |
| PNC EF | 1 | 50 | 0.29 | 0.06 | 5 | 10 | 0.31 | 0.07 | 10 | 6 | 0.31 | 0.07 | 20 | | | |
| | | 60 | 0.27 | 0.05 | | 12 | 0.30 | 0.07 | | 7 | 0.30 | 0.07 | | | | |
| | | 70 | 0.23 | 0.03 | | 14 | 0.28 | 0.06 | | 8 | 0.28 | 0.06 | | | | |
| | | 80 | 0.19 | 0.01 | | 16 | 0.26 | 0.05 | | 9 | 0.27 | 0.05 | | | | |
| | | 90 | 0.13 | -0.01 | | 18 | 0.24 | 0.03 | | 10 | 0.18 | 0.01 | | | | |
| | | 100 | 0.01 | -0.03 | | 20 | 0.08 | -0.02 | | | | | | | | |
| | | 1 | 0.40 | 0.15 | | 1 | 0.39 | 0.14 | | 1 | 0.38 | 0.13 | | 1 | 0.38 | 0.13 |
| | | 10 | 0.37 | 0.13 | | 2 | 0.37 | 0.13 | | 2 | 0.36 | 0.12 | | 2 | 0.35 | 0.11 |
| | | 20 | 0.35 | 0.11 | | 4 | 0.36 | 0.12 | | 3 | 0.35 | 0.11 | | 3 | 0.35 | 0.11 |
| | | 30 | 0.33 | 0.10 | | 6 | 0.35 | 0.11 | | 4 | 0.35 | 0.11 | | 4 | 0.34 | 0.10 |
| | | 40 | 0.33 | 0.09 | | 8 | 0.35 | 0.11 | | 5 | 0.35 | 0.11 | | 5 | 0.32 | 0.09 |
| PNC Language | 1 | 50 | 0.32 | 0.09 | 5 | 10 | 0.35 | 0.11 | 10 | 6 | 0.35 | 0.11 | 20 | | | |
| | | 60 | 0.30 | 0.08 | | 12 | 0.34 | 0.10 | | 7 | 0.34 | 0.10 | | | | |
| | | 70 | 0.26 | 0.06 | | 14 | 0.33 | 0.09 | | 8 | 0.33 | 0.10 | | | | |
| | | 80 | 0.22 | 0.04 | | 16 | 0.31 | 0.09 | | 9 | 0.32 | 0.09 | | | | |
| | | 90 | 0.15 | 0.01 | | 18 | 0.28 | 0.07 | | 10 | 0.20 | 0.03 | | | | |
| | | 100 | 0.01 | -0.01 | | 20 | 0.09 | 0.00 | | | | | | | | |
| | | 1 | 0.13 | -0.11 | | 1 | 0.14 | -0.10 | | 1 | 0.14 | -0.10 | | 1 | 0.14 | -0.09 |
| | | 10 | 0.14 | -0.09 | | 2 | 0.14 | -0.10 | | 2 | 0.15 | -0.08 | | 2 | 0.15 | -0.08 |
| | | 20 | 0.14 | -0.08 | | 4 | 0.15 | -0.08 | | 3 | 0.15 | -0.08 | | 3 | 0.16 | -0.07 |
| | | 30 | 0.14 | -0.08 | | 6 | 0.15 | -0.08 | | 4 | 0.15 | -0.07 | | 4 | 0.16 | -0.07 |
| | | 40 | 0.14 | -0.07 | | 8 | 0.15 | -0.07 | | 5 | 0.16 | -0.07 | | 5 | 0.14 | -0.06 |
| HCPD EF | 1 | 50 | 0.12 | -0.08 | 5 | 10 | 0.15 | -0.07 | 10 | 6 | 0.16 | -0.07 | 20 | | | |
| | | 60 | 0.12 | -0.07 | | 12 | 0.15 | -0.07 | | 7 | 0.15 | -0.07 | | | | |
| | | 70 | 0.10 | -0.06 | | 14 | 0.14 | -0.07 | | 8 | 0.15 | -0.07 | | | | |
| | | 80 | 0.07 | -0.05 | | 16 | 0.13 | -0.07 | | 9 | 0.14 | -0.06 | | | | |
| | | 90 | 0.04 | -0.04 | | 18 | 0.10 | -0.06 | | 10 | 0.07 | -0.04 | | | | |
| | | 100 | 0.00 | -0.03 | | 20 | 0.02 | -0.03 | | | | | | | | |
| | | 1 | 0.18 | -0.10 | | 1 | 0.19 | -0.08 | | 1 | 0.18 | -0.10 | | 1 | 0.16 | -0.11 |
| | | 10 | 0.13 | -0.14 | | 2 | 0.16 | -0.12 | | 2 | 0.14 | -0.14 | | 2 | 0.14 | -0.13 |
| | | 20 | 0.12 | -0.14 | | 4 | 0.14 | -0.14 | | 3 | 0.14 | -0.13 | | 3 | 0.16 | -0.11 |
| | | 30 | 0.12 | -0.13 | | 6 | 0.14 | -0.13 | | 4 | 0.14 | -0.13 | | 4 | 0.18 | -0.08 |
| | | 40 | 0.11 | -0.13 | | 8 | 0.14 | -0.13 | | 5 | 0.15 | -0.11 | | 5 | 0.12 | -0.09 |
| HCPD Language | 1 | 50 | 0.12 | -0.11 | 5 | 10 | 0.15 | -0.11 | 10 | 6 | 0.16 | -0.10 | 20 | | | |
| | | 60 | 0.12 | -0.09 | | 12 | 0.16 | -0.10 | | 7 | 0.17 | -0.09 | | | | |
| | | 70 | 0.10 | -0.07 | | 14 | 0.16 | -0.09 | | 8 | 0.16 | -0.09 | | | | |
| | | 80 | 0.07 | -0.06 | | 16 | 0.13 | -0.09 | | 9 | 0.12 | -0.09 | | | | |
| | | 90 | 0.03 | -0.04 | | 18 | 0.08 | -0.08 | | 10 | 0.05 | -0.05 | | | | |
| | | 100 | 0.00 | -0.03 | | 20 | 0.02 | -0.03 | | | | | | | | |
| | | 1 | 0.19 | -0.01 | | 1 | 0.19 | -0.01 | | 1 | 0.18 | -0.01 | | 1 | 0.17 | -0.02 |
| | | 10 | 0.14 | -0.03 | | 2 | 0.16 | -0.02 | | 2 | 0.15 | -0.03 | | 2 | 0.14 | -0.03 |
| | | 20 | 0.13 | -0.03 | | 4 | 0.14 | -0.03 | | 3 | 0.14 | -0.03 | | 3 | 0.14 | -0.02 |
| | | 30 | 0.13 | -0.03 | | 6 | 0.14 | -0.03 | | 4 | 0.14 | -0.03 | | 4 | 0.12 | -0.03 |
| | | 40 | 0.11 | -0.03 | | 8 | 0.14 | -0.03 | | 5 | 0.14 | -0.02 | | 5 | 0.07 | -0.04 |
| HBN EF | 1 | 50 | 0.11 | -0.03 | 5 | 10 | 0.13 | -0.03 | 10 | 6 | 0.13 | -0.03 | 20 | | | |
| | | 60 | 0.09 | -0.03 | | 12 | 0.13 | -0.03 | | 7 | 0.12 | -0.03 | | | | |
| | | 70 | 0.07 | -0.03 | | 14 | 0.11 | -0.03 | | 8 | 0.11 | -0.03 | | | | |
| | | 80 | 0.05 | -0.02 | | 16 | 0.09 | -0.03 | | 9 | 0.08 | -0.04 | | | | |
| | | 90 | 0.02 | -0.02 | | 18 | 0.05 | -0.03 | | 10 | 0.03 | -0.02 | | | | |
| | | 100 | 0.00 | -0.01 | | 20 | 0.01 | -0.01 | | | | | | | | |
| | | 1 | 0.20 | -0.02 | | 1 | 0.20 | -0.01 | | 1 | 0.20 | -0.01 | | 1 | 0.20 | -0.01 |
| | | 10 | 0.19 | -0.01 | | 2 | 0.21 | 0.00 | | 2 | 0.20 | -0.01 | | 2 | 0.18 | -0.01 |
| | | 20 | 0.17 | -0.02 | | 4 | 0.19 | -0.01 | | 3 | 0.18 | -0.02 | | 3 | 0.19 | -0.01 |
| | | 30 | 0.16 | -0.02 | | 6 | 0.17 | -0.02 | | 4 | 0.18 | -0.01 | | 4 | 0.19 | -0.01 |
| | | 40 | 0.15 | -0.02 | | 8 | 0.18 | -0.02 | | 5 | 0.18 | -0.01 | | 5 | 0.14 | -0.02 |
| HBN Language | 1 | 50 | 0.15 | -0.02 | 5 | 10 | 0.18 | -0.01 | 10 | 6 | 0.19 | -0.01 | 20 | | | |
| | | 60 | 0.13 | -0.02 | | 12 | 0.18 | -0.01 | | 7 | 0.18 | -0.01 | | | | |
| | | 70 | 0.10 | -0.02 | | 14 | 0.17 | -0.01 | | 8 | 0.17 | -0.01 | | | | |
| | | 80 | 0.08 | -0.02 | | 16 | 0.14 | -0.02 | | 9 | 0.14 | -0.01 | | | | |
| | | 90 | 0.04 | -0.01 | | 18 | 0.10 | -0.02 | | 10 | 0.06 | -0.02 | | | | |
| | | 100 | 0.00 | -0.01 | | 20 | 0.02 | -0.01 | | | | | | | | |

**Table S3. Within-dataset cognitive phenotype predictions across 1%, 5%, 10%, and 20% of features.** For the percentile analysis, features were grouped into one hundred non-overlapping subsets, each representing 1% of the total features. For the ventile analysis, features were grouped into twenty non-overlapping subsets, each representing 5% of the total features. For the decile analysis, features were grouped into twenty non-overlapping subsets, each representing 5% of the total features. For the quintile analysis, features were divided into five non-overlapping subsets, each representing 20% of the total features.

| Train Phenotype | Test Phenotype | Percent Edges Used | Percentile | Performance | Percent Edges Used | Ventile | Performance | Percent Edges Used | Decile | Performance | Percent Edges Used | Quintile | Performance |
|---|---|---|---|---|---|---|---|---|---|---|---|---|---|
| HBN EF (n = 1110) | PNC EF (n = 1291) | 1 | 1 | r = 0.28 | 5 | 1 | r = 0.29 | 10 | 1 | r = 0.29 | 20 | 1 | r = 0.28 |
| | | | 5 | r = 0.28 | | 2 | r = 0.29 | | 2 | r = 0.27 | | 2 | r = 0.26 |
| | | | 10 | r = 0.27 | | 4 | r = 0.25 | | 3 | r = 0.26 | | 3 | r = 0.25 |
| | | | 20 | r = 0.26 | | 6 | r = 0.24 | | 4 | r = 0.25 | | 4 | r = 0.22 |
| | | | 30 | r = 0.22 | | 8 | r = 0.22 | | 5 | r = 0.24 | | 5 | r = 0.13 |
| | | | 40 | r = 0.20 | | 10 | r = 0.22 | | 6 | r = 0.25 | | | |
| | | | 50 | r = 0.20 | | 12 | r = 0.26 | | 7 | r = 0.21 | | | |
| | | | 60 | r = 0.22 | | 14 | r = 0.19 | | 8 | r = 0.20 | | | |
| | | | 70 | r = 0.12 | | 16 | r = 0.14 | | 9 | r = 0.14 | | | |
| | | | 80 | r = 0.04 | | 18 | r = 0.14 | | 10 | r = 0.05 | | | |
| | | | 90 | r = 0.09 | | 20 | r = -0.02 | | | | | | |
| | | | 100 | r = 0.10 | | | | | | | | | |
| HCDP EF (n = 428) | | 1 | 1 | r = 0.27 | 5 | 1 | r = 0.28 | 10 | 1 | r = 0.29 | 20 | 1 | r = 0.29 |
| | | | 5 | r = 0.28 | | 2 | r = 0.30 | | 2 | r = 0.29 | | 2 | r = 0.29 |
| | | | 10 | r = 0.28 | | 4 | r = 0.29 | | 3 | r = 0.29 | | 3 | r = 0.29 |
| | | | 20 | r = 0.25 | | 6 | r = 0.30 | | 4 | r = 0.29 | | 4 | r = 0.26 |
| | | | 30 | r = 0.27 | | 8 | r = 0.28 | | 5 | r = 0.29 | | 5 | r = 0.22 |
| | | | 40 | r = 0.22 | | 10 | r = 0.29 | | 6 | r = 0.27 | | | |
| | | | 50 | r = 0.27 | | 12 | r = 0.28 | | 7 | r = 0.26 | | | |
| | | | 60 | r = 0.20 | | 14 | r = 0.24 | | 8 | r = 0.23 | | | |
| | | | 70 | r = 0.15 | | 16 | r = 0.18 | | 9 | r = 0.21 | | | |
| | | | 80 | r = 0.05 | | 18 | r = 0.17 | | 10 | r = 0.14 | | | |
| | | | 90 | r = 0.07 | | 20 | r = 0.08 | | | | | | |
| | | | 100 | r = 0.05 | | | | | | | | | |
| HBN Language (n = 1110) | PNC Language (n = 1291) | 1 | 1 | r = 0.27 | 5 | 1 | r = 0.28 | 10 | 1 | r = 0.27 | 20 | 1 | r = 0.26 |
| | | | 5 | r = 0.24 | | 2 | r = 0.25 | | 2 | r = 0.22 | | 2 | r = 0.20 |
| | | | 10 | r = 0.17 | | 4 | r = 0.22 | | 3 | r = 0.20 | | 3 | r = 0.23 |
| | | | 20 | r = 0.19 | | 6 | r = 0.18 | | 4 | r = 0.19 | | 4 | r = 0.20 |
| | | | 30 | r = 0.08 | | 8 | r = 0.19 | | 5 | r = 0.23 | | 5 | r = 0.20 |
| | | | 40 | r = 0.07 | | 10 | r = 0.25 | | 6 | r = 0.20 | | | |
| | | | 50 | r = 0.09 | | 12 | r = 0.16 | | 7 | r = 0.22 | | | |
| | | | 60 | r = 0.12 | | 14 | r = 0.21 | | 8 | r = 0.13 | | | |
| | | | 70 | r = 0.18 | | 16 | r = 0.12 | | 9 | r = 0.16 | | | |
| | | | 80 | r = 0.08 | | 18 | r = 0.09 | | 10 | r = 0.13 | | | |
| | | | 90 | r = 0.04 | | 20 | r = 0.09 | | | | | | |
| | | | 100 | r = -0.07 | | | | | | | | | |
| HCDP Language (n = 428) | | 1 | 1 | r = 0.30 | 5 | 1 | r = 0.31 | 10 | 1 | r = 0.32 | 20 | 1 | r = 0.32 |
| | | | 5 | r = 0.30 | | 2 | r = 0.31 | | 2 | r = 0.33 | | 2 | r = 0.34 |
| | | | 10 | r = 0.27 | | 4 | r = 0.33 | | 3 | r = 0.33 | | 3 | r = 0.33 |
| | | | 20 | r = 0.30 | | 6 | r = 0.30 | | 4 | r = 0.34 | | 4 | r = 0.31 |
| | | | 30 | r = 0.20 | | 8 | r = 0.33 | | 5 | r = 0.33 | | 5 | r = 0.26 |
| | | | 40 | r = 0.28 | | 10 | r = 0.31 | | 6 | r = 0.29 | | | |
| | | | 50 | r = 0.24 | | 12 | r = 0.27 | | 7 | r = 0.29 | | | |
| | | | 60 | r = 0.13 | | 14 | r = 0.26 | | 8 | r = 0.30 | | | |
| | | | 70 | r = 0.20 | | 16 | r = 0.23 | | 9 | r = 0.25 | | | |
| | | | 80 | r = 0.06 | | 18 | r = 0.21 | | 10 | r = 0.16 | | | |
| | | | 90 | r = 0.11 | | 20 | r = 0.09 | | | | | | |
| | | | 100 | r = 0.05 | | | | | | | | | |

**Table S4. Cross-dataset performances testing in PNC.** Decile-based predictions with performance r >= 0.05 reach one-tailed significance.

| Train Phenotype | Test Phenotype | Percent Edges Used | Percentile | Performance | Percent Edges Used | Ventile | Performance | Percent Edges Used | Decile | Performance | Percent Edges Used | Quintile | Performance |
|---|---|---|---|---|---|---|---|---|---|---|---|---|---|
| HBN EF (n = 1110) | HCPD EF (n = 428) | 1 | 1 | r = 0.12 | 5 | 1 | r = 0.13 | 10 | 1 | r = 0.13 | 20 | 1 | r = 0.14 |
| | | | 5 | r = 0.14 | | 2 | r = 0.14 | | 2 | r = 0.14 | | 2 | r = 0.12 |
| | | | 10 | r = 0.13 | | 4 | r = 0.14 | | 3 | r = 0.12 | | 3 | r = 0.13 |
| | | | 20 | r = 0.15 | | 6 | r = 0.10 | | 4 | r = 0.11 | | 4 | r = 0.11 |
| | | | 30 | r = 0.06 | | 8 | r = 0.10 | | 5 | r = 0.13 | | 5 | r = 0.04 |
| | | | 40 | r = 0.03 | | 10 | r = 0.09 | | 6 | r = 0.13 | | | |
| | | | 50 | r = 0.08 | | 12 | r = 0.13 | | 7 | r = 0.12 | | | |
| | | | 60 | r = 0.10 | | 14 | r = 0.10 | | 8 | r = 0.09 | | | |
| | | | 70 | r = 0.11 | | 16 | r = 0.09 | | 9 | r = 0.01 | | | |
| | | | 80 | r = 0.05 | | 18 | r = 0.01 | | 10 | r = 0.06 | | | |
| | | | 90 | r = 0.04 | | 20 | r = 0.11 | | | | | | |
| | | | 100 | r = -0.12 | | | | | | | | | |
| PNC EF (n = 1291) | | 1 | 1 | r = 0.14 | 5 | 1 | r = 0.14 | 10 | 1 | r = 0.14 | 20 | 1 | r = 0.14 |
| | | | 5 | r = 0.15 | | 2 | r = 0.14 | | 2 | r = 0.14 | | 2 | r = 0.13 |
| | | | 10 | r = 0.11 | | 4 | r = 0.14 | | 3 | r = 0.13 | | 3 | r = 0.13 |
| | | | 20 | r = 0.13 | | 6 | r = 0.13 | | 4 | r = 0.13 | | 4 | r = 0.13 |
| | | | 30 | r = 0.15 | | 8 | r = 0.13 | | 5 | r = 0.12 | | 5 | r = 0.13 |
| | | | 40 | r = 0.13 | | 10 | r = 0.13 | | 6 | r = 0.14 | | | |
| | | | 50 | r = 0.15 | | 12 | r = 0.14 | | 7 | r = 0.12 | | | |
| | | | 60 | r = 0.12 | | 14 | r = 0.10 | | 8 | r = 0.12 | | | |
| | | | 70 | r = 0.08 | | 16 | r = 0.13 | | 9 | r = 0.13 | | | |
| | | | 80 | r = 0.08 | | 18 | r = 0.16 | | 10 | r = 0.05 | | | |
| | | | 90 | r = 0.09 | | 20 | r = 0.06 | | | | | | |
| | | | 100 | r = -0.01 | | | | | | | | | |
| HBN Language (n = 1110) | HCPD Language (n = 428) | 1 | 1 | r = 0.09 | 5 | 1 | r = 0.12 | 10 | 1 | r = 0.12 | 20 | 1 | r = 0.13 |
| | | | 5 | r = 0.14 | | 2 | r = 0.12 | | 2 | r = 0.13 | | 2 | r = 0.11 |
| | | | 10 | r = 0.08 | | 4 | r = 0.12 | | 3 | r = 0.09 | | 3 | r = 0.13 |
| | | | 20 | r = 0.11 | | 6 | r = 0.06 | | 4 | r = 0.12 | | 4 | r = 0.13 |
| | | | 30 | r = 0.04 | | 8 | r = 0.11 | | 5 | r = 0.13 | | 5 | r = 0.17 |
| | | | 40 | r = 0.10 | | 10 | r = 0.13 | | 6 | r = 0.12 | | | |
| | | | 50 | r = 0.07 | | 12 | r = 0.08 | | 7 | r = 0.12 | | | |
| | | | 60 | r = 0.09 | | 14 | r = 0.13 | | 8 | r = 0.10 | | | |
| | | | 70 | r = 0.18 | | 16 | r = 0.06 | | 9 | r = 0.08 | | | |
| | | | 80 | r = 0.06 | | 18 | r = 0.05 | | 10 | r = 0.18 | | | |
| | | | 90 | r = -0.02 | | 20 | r = 0.09 | | | | | | |
| | | | 100 | r = -0.10 | | | | | | | | | |
| PNC Language (n = 1291) | | 1 | 1 | r = 0.15 | 5 | 1 | r = 0.13 | 10 | 1 | r = 0.14 | 20 | 1 | r = 0.13 |
| | | | 5 | r = 0.13 | | 2 | r = 0.14 | | 2 | r = 0.13 | | 2 | r = 0.12 |
| | | | 10 | r = 0.13 | | 4 | r = 0.13 | | 3 | r = 0.12 | | 3 | r = 0.12 |
| | | | 20 | r = 0.12 | | 6 | r = 0.12 | | 4 | r = 0.12 | | 4 | r = 0.11 |
| | | | 30 | r = 0.09 | | 8 | r = 0.11 | | 5 | r = 0.11 | | 5 | r = 0.10 |
| | | | 40 | r = 0.06 | | 10 | r = 0.12 | | 6 | r = 0.12 | | | |
| | | | 50 | r = 0.11 | | 12 | r = 0.11 | | 7 | r = 0.11 | | | |
| | | | 60 | r = 0.12 | | 14 | r = 0.11 | | 8 | r = 0.11 | | | |
| | | | 70 | r = 0.02 | | 16 | r = 0.12 | | 9 | r = 0.07 | | | |
| | | | 80 | r = 0.04 | | 18 | r = 0.00 | | 10 | r = 0.12 | | | |
| | | | 90 | r = 0.02 | | 20 | r = 0.07 | | | | | | |
| | | | 100 | r = 0.04 | | | | | | | | | |

**Table S5. Cross-dataset performances testing in HCPD.** Decile-based predictions with performance r >= 0.08 reach one-tailed significance.

| Train Phenotype | Test Phenotype | Percent Edges Used | Percentile | Performance | Percent Edges Used | Ventile | Performance | Percent Edges Used | Decile | Performance | Percent Edges Used | Quintile | Performance |
|---|---|---|---|---|---|---|---|---|---|---|---|---|---|
| PNC EF (n = 1291) | HBN EF (n = 1110) | 1 | 1 | r = 0.11 | 5 | 1 | r = 0.11 | 10 | 1 | r = 0.11 | 20 | 1 | r = 0.11 |
| | | | 5 | r = 0.09 | | 2 | r = 0.10 | | 2 | r = 0.10 | | 2 | r = 0.09 |
| | | | 10 | r = 0.09 | | 4 | r = 0.10 | | 3 | r = 0.10 | | 3 | r = 0.09 |
| | | | 20 | r = 0.12 | | 6 | r = 0.10 | | 4 | r = 0.08 | | 4 | r = 0.11 |
| | | | 30 | r = 0.10 | | 8 | r = 0.08 | | 5 | r = 0.09 | | 5 | r = 0.04 |
| | | | 40 | r = 0.08 | | 10 | r = 0.09 | | 6 | r = 0.10 | | | |
| | | | 50 | r = 0.08 | | 12 | r = 0.10 | | 7 | r = 0.11 | | | |
| | | | 60 | r = 0.09 | | 14 | r = 0.07 | | 8 | r = 0.11 | | | |
| | | | 70 | r = 0.06 | | 16 | r = 0.10 | | 9 | r = 0.06 | | | |
| | | | 80 | r = 0.02 | | 18 | r = 0.05 | | 10 | r = -0.02 | | | |
| | | | 90 | r = 0.06 | | 20 | r = -0.03 | | | | | | |
| | | | 100 | r = -0.05 | | | | | | | | | |
| HCDP EF (n = 428) | | 1 | 1 | r = 0.08 | 5 | 1 | r = 0.08 | 10 | 1 | r = 0.08 | 20 | 1 | r = 0.09 |
| | | | 5 | r = 0.07 | | 2 | r = 0.08 | | 2 | r = 0.09 | | 2 | r = 0.10 |
| | | | 10 | r = 0.06 | | 4 | r = 0.09 | | 3 | r = 0.09 | | 3 | r = 0.10 |
| | | | 20 | r = 0.05 | | 6 | r = 0.09 | | 4 | r = 0.11 | | 4 | r = 0.10 |
| | | | 30 | r = 0.08 | | 8 | r = 0.10 | | 5 | r = 0.10 | | 5 | r = 0.06 |
| | | | 40 | r = 0.08 | | 10 | r = 0.11 | | 6 | r = 0.10 | | | |
| | | | 50 | r = 0.09 | | 12 | r = 0.09 | | 7 | r = 0.08 | | | |
| | | | 60 | r = 0.02 | | 14 | r = 0.05 | | 8 | r = 0.10 | | | |
| | | | 70 | r = 0.04 | | 16 | r = 0.09 | | 9 | r = 0.04 | | | |
| | | | 80 | r = -0.02 | | 18 | r = 0.02 | | 10 | r = 0.06 | | | |
| | | | 90 | r = 0.00 | | 20 | r = 0.02 | | | | | | |
| | | | 100 | r = -0.01 | | | | | | | | | |
| PNC Language (n = 1291) | HBN Language (n = 1110) | 1 | 1 | r = 0.09 | 5 | 1 | r = 0.07 | 10 | 1 | r = 0.07 | 20 | 1 | r = 0.06 |
| | | | 5 | r = 0.03 | | 2 | r = 0.06 | | 2 | r = 0.06 | | 2 | r = 0.06 |
| | | | 10 | r = 0.05 | | 4 | r = 0.06 | | 3 | r = 0.05 | | 3 | r = 0.05 |
| | | | 20 | r = 0.02 | | 6 | r = 0.05 | | 4 | r = 0.06 | | 4 | r = 0.05 |
| | | | 30 | r = 0.02 | | 8 | r = 0.05 | | 5 | r = 0.04 | | 5 | r = 0.01 |
| | | | 40 | r = 0.05 | | 10 | r = 0.03 | | 6 | r = 0.05 | | | |
| | | | 50 | r = 0.05 | | 12 | r = 0.05 | | 7 | r = 0.05 | | | |
| | | | 60 | r = 0.06 | | 14 | r = 0.05 | | 8 | r = 0.04 | | | |
| | | | 70 | r = -0.01 | | 16 | r = 0.03 | | 9 | r = 0.03 | | | |
| | | | 80 | r = -0.01 | | 18 | r = 0.03 | | 10 | r = -0.01 | | | |
| | | | 90 | r = -0.02 | | 20 | r = -0.01 | | | | | | |
| | | | 100 | r = -0.01 | | | | | | | | | |
| HCDP Language (n = 428) | | 1 | 1 | r = 0.06 | 5 | 1 | r = 0.08 | 10 | 1 | r = 0.07 | 20 | 1 | r = 0.07 |
| | | | 5 | r = 0.08 | | 2 | r = 0.05 | | 2 | r = 0.06 | | 2 | r = 0.06 |
| | | | 10 | r = 0.06 | | 4 | r = 0.06 | | 3 | r = 0.06 | | 3 | r = 0.05 |
| | | | 20 | r = 0.05 | | 6 | r = 0.06 | | 4 | r = 0.07 | | 4 | r = 0.05 |
| | | | 30 | r = 0.06 | | 8 | r = 0.05 | | 5 | r = 0.04 | | 5 | r = 0.11 |
| | | | 40 | r = 0.02 | | 10 | r = 0.02 | | 6 | r = 0.05 | | | |
| | | | 50 | r = 0.00 | | 12 | r = 0.06 | | 7 | r = 0.06 | | | |
| | | | 60 | r = -0.01 | | 14 | r = 0.05 | | 8 | r = 0.03 | | | |
| | | | 70 | r = 0.05 | | 16 | r = -0.01 | | 9 | r = 0.11 | | | |
| | | | 80 | r = -0.06 | | 18 | r = 0.03 | | 10 | r = 0.05 | | | |
| | | | 90 | r = 0.02 | | 20 | r = 0.00 | | | | | | |
| | | | 100 | r = 0.01 | | | | | | | | | |

**Table S6. Cross-dataset performances testing in HBN.** Decile-based predictions with performance r >= 0.05 reach one-tailed significance.

| Decile | PNC EF | PNC Language | HCPD EF | HCPD Language | HBN EF | HBN Language |
|---|---|---|---|---|---|---|
| 1 | 3572 | 3624 | 3366 | 3357 | 3401 | 3483 |
| 2 | 3803 | 3747 | 2286 | 2362 | 2765 | 2730 |
| 3 | 3287 | 3328 | 1143 | 1051 | 1390 | 1456 |
| 4 | 2707 | 2531 | 687 | 601 | 806 | 881 |
| 5 | 2087 | 2103 | 439 | 413 | 559 | 603 |
| 6 | 1671 | 1672 | 334 | 302 | 339 | 395 |
| 7 | 1542 | 1533 | 347 | 291 | 335 | 357 |
| 8 | 1235 | 1372 | 470 | 374 | 483 | 497 |
| 9 | 1754 | 1843 | 876 | 843 | 967 | 1046 |
| 10 | 3325 | 3420 | 1425 | 1179 | 1487 | 1655 |

**Table S7. Edge selection across folds for each decile-based model.** Counts show the number of edges that were present in at least five of the ten cross-validation folds.

| Phenotype | Percent of Edges | Decile | Performance | |
|---|---|---|---|---|
| | | | r | q2 |
| PNC EF | 100 | N/A | 0.35 | 0.12 |
| | 10 | 1 | 0.36 | 0.12 |
| | | 2 | 0.34 | 0.11 |
| | | 3 | 0.34 | 0.11 |
| | | 4 | 0.31 | 0.09 |
| | | 5 | 0.30 | 0.08 |
| | | 6 | 0.27 | 0.05 |
| | | 7 | 0.10 | 0.00 |
| | | 8 | -0.01 | 0.00 |
| | | 9 | -0.05 | 0.00 |
| | | 10 | -0.07 | 0.00 |
| PNC Language | 100 | N/A | 0.43 | 0.18 |
| | 10 | 1 | 0.45 | 0.19 |
| | | 2 | 0.44 | 0.19 |
| | | 3 | 0.43 | 0.17 |
| | | 4 | 0.37 | 0.11 |
| | | 5 | 0.36 | 0.11 |
| | | 6 | 0.33 | 0.09 |
| | | 7 | 0.14 | 0.01 |
| | | 8 | 0.01 | 0.00 |
| | | 9 | -0.05 | 0.00 |
| | | 10 | -0.07 | 0.00 |
| HCPD EF | 100 | N/A | 0.07 | -0.12 |
| | 10 | 1 | 0.12 | -0.09 |
| | | 2 | 0.12 | -0.05 |
| | | 3 | 0.08 | 0.00 |
| | | 4 | 0.05 | 0.00 |
| | | 5 | 0.07 | 0.00 |
| | | 6 | 0.07 | 0.00 |
| | | 7 | 0.05 | 0.00 |
| | | 8 | 0.02 | 0.00 |
| | | 9 | 0.01 | 0.00 |
| | | 10 | 0.00 | 0.00 |
| HCPD Language | 100 | N/A | 0.08 | -0.19 |
| | 10 | 1 | 0.21 | -0.04 |
| | | 2 | 0.13 | -0.07 |
| | | 3 | 0.06 | -0.02 |
| | | 4 | 0.04 | 0.00 |
| | | 5 | 0.07 | 0.00 |
| | | 6 | 0.05 | 0.00 |
| | | 7 | 0.04 | 0.00 |
| | | 8 | 0.02 | 0.00 |
| | | 9 | 0.01 | 0.00 |
| | | 10 | -0.01 | 0.00 |
| HBN EF | 100 | N/A | 0.13 | 0.00 |
| | 10 | 1 | 0.19 | -0.02 |
| | | 2 | 0.15 | -0.03 |
| | | 3 | 0.14 | 0.01 |
| | | 4 | 0.10 | 0.01 |
| | | 5 | -0.03 | 0.00 |
| | | 6 | -0.05 | 0.00 |
| | | 7 | -0.06 | 0.00 |
| | | 8 | -0.07 | 0.00 |
| | | 9 | -0.08 | 0.00 |
| | | 10 | -0.08 | 0.00 |
| HBN Language | 100 | N/A | 0.17 | 0.02 |
| | 10 | 1 | 0.28 | 0.03 |
| | | 2 | 0.25 | 0.02 |
| | | 3 | 0.20 | 0.03 |
| | | 4 | 0.15 | 0.02 |
| | | 5 | 0.00 | 0.00 |
| | | 6 | -0.04 | 0.00 |
| | | 7 | -0.06 | 0.00 |
| | | 8 | -0.07 | 0.00 |
| | | 9 | -0.08 | 0.00 |
| | | 10 | -0.09 | 0.00 |

**Table S8. Ridge regression cognitive phenotype predictions using all features and decile-based subsets.** To evaluate the robustness of our findings across modeling approaches, we implemented ridge regression using both the full connectome (no feature selection) and each decile subset of features. Deciles were defined based on the univariate correlation between each feature and the target phenotype, with decile 1 containing the most strongly associated features and decile 10 the weakest.

| Comparison | p value |
|---|---|
| All Features vs. Decile 1 | 0.02 |
| All Features vs. Decile 2 | 0.05 |
| All Features vs. Decile 3 | 0.66 |
| All Features vs. Decile 4 | 0.00 |
| All Features vs. Decile 5 | 0.05 |
| Decile 1 vs. Decile 2 | 0.05 |
| Decile 1 vs. Decile 3 | 0.03 |
| Decile 1 vs. Decile 4 | 0.00 |
| Decile 1 vs. Decile 5 | 0.01 |
| Decile 2 vs. Decile 3 | 0.04 |
| Decile 2 vs. Decile 4 | 0.00 |
| Decile 2 vs. Decile 5 | 0.03 |
| Decile 3 vs. Decile 4 | 0.00 |
| Decile 3 vs. Decile 4 | 0.07 |
| Decile 4 vs. Decile 5 | 0.25 |

**Table S9. Paired t-tests for ridge regression comparing decile 1 to other deciles across executive function and language models.** To assess whether features from a given feature set yielded significantly better prediction performance than other feature sets, we conducted paired t-tests. The comparison was performed across the six primary models (executive function and language phenotypes for PNC, HBN, and HCPD). Each p-value reflects a paired t-test comparing the prediction accuracy (e.g., Pearson's r) of a given feature set versus another feature set across the six models.

| Phenotype | Percent of Edges | Percentile | Performance | | Percent of Edges | Ventile | Performance | | Percent of Edges | Decile | Performance | | Percent of Edges | Quintile | Performance | |
|---|---|---|---|---|---|---|---|---|---|---|---|---|---|---|---|---|
| | | | r | q2 | | | r | q2 | | | r | q2 | | | r | q2 |
| | | 1 | 0.30 | 0.03 | | 1 | 0.33 | 0.07 | | 1 | 0.34 | 0.08 | | 1 | 0.34 | 0.08 |
| | | 10 | 0.20 | -0.05 | | 2 | 0.31 | 0.06 | | 2 | 0.31 | 0.05 | | 2 | 0.32 | 0.07 |
| | | 20 | 0.16 | -0.08 | | 4 | 0.27 | 0.02 | | 3 | 0.29 | 0.05 | | 3 | 0.28 | 0.03 |
| | | 30 | 0.14 | -0.09 | | 6 | 0.25 | 0.00 | | 4 | 0.28 | 0.03 | | 4 | 0.20 | -0.05 |
| | | 40 | 0.11 | -0.09 | | 8 | 0.22 | -0.03 | | 5 | 0.25 | 0.00 | | 5 | 0.07 | -0.07 |
| PNC EF | 1 | 50 | 0.09 | -0.08 | 5 | 10 | 0.19 | -0.06 | 10 | 6 | 0.21 | -0.04 | 20 | | | |
| | | 60 | 0.08 | -0.07 | | 12 | 0.16 | -0.08 | | 7 | 0.18 | -0.07 | | | | |
| | | 70 | 0.05 | -0.06 | | 14 | 0.12 | -0.09 | | 8 | 0.13 | -0.09 | | | | |
| | | 80 | 0.04 | -0.05 | | 16 | 0.09 | -0.07 | | 9 | 0.08 | -0.08 | | | | |
| | | 90 | 0.02 | -0.04 | | 18 | 0.05 | -0.05 | | 10 | 0.02 | -0.04 | | | | |
| | | 100 | 0.01 | -0.04 | | 20 | 0.01 | -0.04 | | | | | | | | |
| | | 1 | 0.46 | 0.18 | | 1 | 0.51 | 0.24 | | 1 | 0.50 | 0.23 | | 1 | 0.49 | 0.22 |
| | | 10 | 0.29 | 0.01 | | 2 | 0.44 | 0.18 | | 2 | 0.43 | 0.16 | | 2 | 0.43 | 0.17 |
| | | 20 | 0.22 | -0.06 | | 4 | 0.38 | 0.11 | | 3 | 0.41 | 0.14 | | 3 | 0.38 | 0.12 |
| | | 30 | 0.19 | -0.09 | | 6 | 0.34 | 0.08 | | 4 | 0.37 | 0.11 | | 4 | 0.28 | 0.01 |
| | | 40 | 0.15 | -0.11 | | 8 | 0.29 | 0.03 | | 5 | 0.33 | 0.07 | | 5 | 0.10 | -0.10 |
| PNC Language | 1 | 50 | 0.13 | -0.11 | 5 | 10 | 0.25 | -0.03 | 10 | 6 | 0.29 | 0.02 | 20 | | | |
| | | 60 | 0.10 | -0.09 | | 12 | 0.22 | -0.06 | | 7 | 0.24 | -0.04 | | | | |
| | | 70 | 0.07 | -0.07 | | 14 | 0.16 | -0.11 | | 8 | 0.19 | -0.09 | | | | |
| | | 80 | 0.05 | -0.04 | | 16 | 0.13 | -0.10 | | 9 | 0.11 | -0.10 | | | | |
| | | 90 | 0.02 | -0.03 | | 18 | 0.07 | -0.06 | | 10 | 0.04 | -0.03 | | | | |
| | | 100 | 0.01 | -0.02 | | 20 | 0.01 | -0.02 | | | | | | | | |

**Table S10. Partial correlation CPM predictions across 1%, 5%, 10%, and 20% of features.** To assess whether global effects in Pearson-based functional connectivity might influence results, we repeated our decile-based predictive modeling using partial correlation connectomes. These were derived from concatenated time series across resting-state, n-back, and emotion fMRI runs in the 656 PNC participants with complete data. Partial correlation removes shared variance between nodes and better isolates direct connections, offering a complementary perspective to standard Pearson correlation. This figure shows the prediction performance (Pearson's r) for executive function and language phenotypes when CPM models are trained on the top 1%, 5%, 10%, and 20% of partial correlation features, as ranked by their univariate correlation with the phenotype.

| Phenotype | Percent of Edges | Decile | Performance | |
|---|---|---|---|---|
| | | | r | q2 |
| PNC EF | 100 | N/A | 0.21 | 0.04 |
| | 10 | 1 | 0.29 | 0.07 |
| | | 2 | 0.24 | 0.05 |
| | | 3 | 0.20 | 0.02 |
| | | 4 | 0.01 | 0.00 |
| | | 5 | -0.11 | 0.00 |
| | | 6 | -0.12 | 0.00 |
| | | 7 | -0.12 | 0.00 |
| | | 8 | -0.12 | 0.00 |
| | | 9 | -0.12 | 0.00 |
| | | 10 | -0.12 | 0.00 |
| PNC Language | 100 | N/A | 0.36 | 0.11 |
| | 10 | 1 | 0.48 | 0.22 |
| | | 2 | 0.37 | 0.13 |
| | | 3 | 0.32 | 0.10 |
| | | 4 | 0.10 | 0.01 |
| | | 5 | -0.10 | 0.00 |
| | | 6 | -0.11 | 0.00 |
| | | 7 | -0.11 | 0.00 |
| | | 8 | -0.11 | 0.00 |
| | | 9 | -0.11 | 0.00 |
| | | 10 | -0.11 | 0.00 |

**Table S11. Partial correlation ridge regression predictions.** Analogous to above, predictive modeling was repeated using partial correlation connectomes with ridge regression. These were derived from concatenated time series across resting-state, n-back, and emotion fMRI runs in the 656 PNC participants with complete data.

| Phenotpye | Percent Edges Used | Percentile | Performance | | Percent Edges Used | Ventile | Performance | | Percent Edges Used | Decile | Performance | | Percent Edges Used | Quintile | Performance | |
|---|---|---|---|---|---|---|---|---|---|---|---|---|---|---|---|---|
| | | | r | q2 | | | r | q2 | | | r | q2 | | | r | q2 |
| Learning Problems | 1 | 1 | 0.18 | -0.04 | 5 | 1 | 0.18 | -0.03 | 10 | 1 | 0.18 | -0.04 | 20 | 1 | 0.18 | -0.04 |
| | | 10 | 0.17 | -0.04 | | 2 | 0.17 | -0.04 | | 2 | 0.17 | -0.04 | | 2 | 0.15 | -0.05 |
| | | 20 | 0.15 | -0.05 | | 4 | 0.17 | -0.04 | | 3 | 0.15 | -0.05 | | 3 | 0.14 | -0.06 |
| | | 30 | 0.13 | -0.05 | | 6 | 0.15 | -0.05 | | 4 | 0.14 | -0.06 | | 4 | 0.13 | -0.06 |
| | | 40 | 0.12 | -0.06 | | 8 | 0.14 | -0.06 | | 5 | 0.14 | -0.06 | | 5 | 0.08 | -0.06 |
| | | 50 | 0.11 | -0.06 | | 10 | 0.14 | -0.06 | | 6 | 0.14 | -0.06 | | | | |
| | | 60 | 0.10 | -0.05 | | 12 | 0.14 | -0.05 | | 7 | 0.13 | -0.06 | | | | |
| | | 70 | 0.08 | -0.05 | | 14 | 0.12 | -0.06 | | 8 | 0.12 | -0.06 | | | | |
| | | 80 | 0.05 | -0.04 | | 16 | 0.10 | -0.06 | | 9 | 0.08 | -0.06 | | | | |
| | | 90 | 0.02 | -0.03 | | 18 | 0.05 | -0.05 | | 10 | 0.03 | -0.03 | | | | |
| | | 100 | 0.00 | -0.02 | | 20 | 0.01 | -0.02 | | | | | | | | |
| Internet Addiction | 1 | 1 | 0.15 | -0.08 | 5 | 1 | 0.14 | -0.08 | 10 | 1 | 0.14 | -0.07 | 20 | 1 | 0.14 | -0.07 |
| | | 10 | 0.13 | -0.07 | | 2 | 0.14 | -0.07 | | 2 | 0.13 | -0.07 | | 2 | 0.12 | -0.07 |
| | | 20 | 0.11 | -0.07 | | 4 | 0.12 | -0.07 | | 3 | 0.12 | -0.07 | | 3 | 0.13 | -0.06 |
| | | 30 | 0.10 | -0.07 | | 6 | 0.11 | -0.07 | | 4 | 0.12 | -0.07 | | 4 | 0.13 | -0.05 |
| | | 40 | 0.10 | -0.06 | | 8 | 0.12 | -0.07 | | 5 | 0.13 | -0.06 | | 5 | 0.09 | -0.05 |
| | | 50 | 0.11 | -0.05 | | 10 | 0.13 | -0.06 | | 6 | 0.13 | -0.05 | | | | |
| | | 60 | 0.10 | -0.05 | | 12 | 0.13 | -0.05 | | 7 | 0.13 | -0.05 | | | | |
| | | 70 | 0.08 | -0.04 | | 14 | 0.12 | -0.05 | | 8 | 0.12 | -0.05 | | | | |
| | | 80 | 0.05 | -0.03 | | 16 | 0.10 | -0.05 | | 9 | 0.09 | -0.05 | | | | |
| | | 90 | 0.03 | -0.02 | | 18 | 0.07 | -0.04 | | 10 | 0.04 | -0.03 | | | | |
| | | 100 | 0.00 | -0.02 | | 20 | 0.02 | -0.02 | | | | | | | | |
| Parent/Child Internet Addiction | 1 | 1 | 0.13 | -0.08 | 5 | 1 | 0.14 | -0.05 | 10 | 1 | 0.15 | -0.04 | 20 | 1 | 0.15 | -0.04 |
| | | 10 | 0.15 | -0.03 | | 2 | 0.16 | -0.03 | | 2 | 0.15 | -0.03 | | 2 | 0.14 | -0.04 |
| | | 20 | 0.14 | -0.04 | | 4 | 0.15 | -0.03 | | 3 | 0.14 | -0.04 | | 3 | 0.16 | -0.03 |
| | | 30 | 0.13 | -0.04 | | 6 | 0.14 | -0.04 | | 4 | 0.14 | -0.04 | | 4 | 0.15 | -0.03 |
| | | 40 | 0.13 | -0.04 | | 8 | 0.14 | -0.04 | | 5 | 0.15 | -0.03 | | 5 | 0.10 | -0.04 |
| | | 50 | 0.13 | -0.03 | | 10 | 0.15 | -0.03 | | 6 | 0.16 | -0.03 | | | | |
| | | 60 | 0.12 | -0.03 | | 12 | 0.15 | -0.03 | | 7 | 0.15 | -0.03 | | | | |
| | | 70 | 0.10 | -0.03 | | 14 | 0.14 | -0.03 | | 8 | 0.13 | -0.04 | | | | |
| | | 80 | 0.06 | -0.03 | | 16 | 0.11 | -0.04 | | 9 | 0.10 | -0.04 | | | | |
| | | 90 | 0.03 | -0.02 | | 18 | 0.08 | -0.04 | | 10 | 0.05 | -0.03 | | | | |
| | | 100 | -0.01 | -0.02 | | 20 | 0.02 | -0.02 | | | | | | | | |
| Separation Anxiety | 1 | 1 | 0.18 | -0.07 | 5 | 1 | 0.19 | -0.06 | 10 | 1 | 0.19 | -0.06 | 20 | 1 | 0.20 | -0.05 |
| | | 10 | 0.18 | -0.05 | | 2 | 0.19 | -0.05 | | 2 | 0.20 | -0.04 | | 2 | 0.20 | -0.04 |
| | | 20 | 0.18 | -0.05 | | 4 | 0.20 | -0.04 | | 3 | 0.20 | -0.04 | | 3 | 0.19 | -0.05 |
| | | 30 | 0.17 | -0.05 | | 6 | 0.20 | -0.04 | | 4 | 0.20 | -0.04 | | 4 | 0.15 | -0.07 |
| | | 40 | 0.15 | -0.05 | | 8 | 0.19 | -0.04 | | 5 | 0.19 | -0.05 | | 5 | 0.07 | -0.08 |
| | | 50 | 0.13 | -0.06 | | 10 | 0.18 | -0.05 | | 6 | 0.18 | -0.05 | | | | |
| | | 60 | 0.11 | -0.05 | | 12 | 0.17 | -0.06 | | 7 | 0.16 | -0.07 | | | | |
| | | 70 | 0.08 | -0.05 | | 14 | 0.14 | -0.07 | | 8 | 0.12 | -0.08 | | | | |
| | | 80 | 0.05 | -0.04 | | 16 | 0.09 | -0.08 | | 9 | 0.07 | -0.09 | | | | |
| | | 90 | 0.02 | -0.03 | | 18 | 0.05 | -0.06 | | 10 | 0.03 | -0.04 | | | | |
| | | 100 | 0.01 | -0.02 | | 20 | 0.02 | -0.02 | | | | | | | | |
| Social Communication | 1 | 1 | 0.14 | -0.03 | 5 | 1 | 0.14 | -0.03 | 10 | 1 | 0.14 | -0.03 | 20 | 1 | 0.14 | -0.02 |
| | | 10 | 0.14 | -0.02 | | 2 | 0.14 | -0.03 | | 2 | 0.15 | -0.02 | | 2 | 0.15 | -0.02 |
| | | 20 | 0.14 | -0.02 | | 4 | 0.15 | -0.02 | | 3 | 0.15 | -0.02 | | 3 | 0.16 | -0.02 |
| | | 30 | 0.14 | -0.02 | | 6 | 0.15 | -0.02 | | 4 | 0.16 | -0.02 | | 4 | 0.15 | -0.02 |
| | | 40 | 0.14 | -0.02 | | 8 | 0.16 | -0.02 | | 5 | 0.16 | -0.02 | | 5 | 0.11 | -0.04 |
| | | 50 | 0.14 | -0.02 | | 10 | 0.16 | -0.02 | | 6 | 0.16 | -0.02 | | | | |
| | | 60 | 0.13 | -0.03 | | 12 | 0.15 | -0.02 | | 7 | 0.15 | -0.02 | | | | |
| | | 70 | 0.10 | -0.03 | | 14 | 0.14 | -0.02 | | 8 | 0.13 | -0.03 | | | | |
| | | 80 | 0.07 | -0.04 | | 16 | 0.12 | -0.03 | | 9 | 0.11 | -0.04 | | | | |
| | | 90 | 0.03 | -0.03 | | 18 | 0.08 | -0.04 | | 10 | 0.05 | -0.04 | | | | |
| | | 100 | 0.00 | -0.03 | | 20 | 0.02 | -0.03 | | | | | | | | |
| Hyperactivity | 1 | 1 | 0.16 | -0.05 | 5 | 1 | 0.15 | -0.05 | 10 | 1 | 0.14 | -0.06 | 20 | 1 | 0.14 | -0.06 |
| | | 10 | 0.12 | -0.07 | | 2 | 0.13 | -0.07 | | 2 | 0.13 | -0.06 | | 2 | 0.12 | -0.06 |
| | | 20 | 0.12 | -0.06 | | 4 | 0.13 | -0.06 | | 3 | 0.12 | -0.06 | | 3 | 0.12 | -0.06 |
| | | 30 | 0.11 | -0.06 | | 6 | 0.12 | -0.06 | | 4 | 0.12 | -0.06 | | 4 | 0.10 | -0.07 |
| | | 40 | 0.10 | -0.06 | | 8 | 0.12 | -0.06 | | 5 | 0.12 | -0.06 | | 5 | 0.07 | -0.07 |
| | | 50 | 0.09 | -0.06 | | 10 | 0.11 | -0.06 | | 6 | 0.11 | -0.07 | | | | |
| | | 60 | 0.08 | -0.06 | | 12 | 0.10 | -0.07 | | 7 | 0.10 | -0.07 | | | | |
| | | 70 | 0.06 | -0.05 | | 14 | 0.09 | -0.07 | | 8 | 0.09 | -0.08 | | | | |
| | | 80 | 0.04 | -0.04 | | 16 | 0.07 | -0.07 | | 9 | 0.07 | -0.07 | | | | |
| | | 90 | 0.03 | -0.02 | | 18 | 0.05 | -0.05 | | 10 | 0.03 | -0.03 | | | | |
| | | 100 | 0.01 | -0.02 | | 20 | 0.01 | -0.02 | | | | | | | | |

**Table S12. Psychiatric and developmental predictions across 1%, 5%, 10%, and 20% of features.**

| Dataset | Phenotpye | Percent Edges Used | Percentile | Performance | | | Percent Edges Used | Percentile | Performance | | | Percent Edges Used | Percentile | | Performance | | Percent Edges Used | Percentile | Performance | | |
|---|---|---|---|---|---|---|---|---|---|---|---|---|---|---|---|---|---|---|---|---|---|
| | | | | AUC | r | q2 | | | AUC | r | q2 | | | AUC | r | q2 | | | AUC | r | q2 |
| HBN | Sex | 1 | 1 | 0.68 | | | 5 | 1 | 0.68 | | | 10 | 1 | 0.68 | | | 20 | 1 | 0.68 | | |
| | | | 10 | 0.67 | | | | 2 | 0.67 | | | | 2 | 0.68 | | | | 2 | 0.68 | | |
| | | | 20 | 0.67 | | | | 4 | 0.68 | | | | 3 | 0.68 | | | | 3 | 0.67 | | |
| | | | 30 | 0.67 | | | | 6 | 0.68 | | | | 4 | 0.68 | | | | 4 | 0.66 | | |
| | | | 40 | 0.66 | | | | 8 | 0.68 | | | | 5 | 0.67 | | | | 5 | 0.63 | | |
| | | | 50 | 0.64 | | | | 10 | 0.67 | | | | 6 | 0.67 | | | | | | | |
| | | | 60 | 0.63 | | | | 12 | 0.66 | | | | 7 | 0.66 | | | | | | | |
| | | | 70 | 0.61 | | | | 14 | 0.65 | | | | 8 | 0.66 | | | | | | | |
| | | | 80 | 0.59 | | | | 16 | 0.64 | | | | 9 | 0.64 | | | | | | | |
| | | | 90 | 0.54 | | | | 18 | 0.60 | | | | 10 | 0.56 | | | | | | | |
| | | | 100 | 0.45 | | | | 20 | 0.52 | | | | | | | | | | | | |
| | Age | 1 | 1 | | 0.55 | 0.29 | 5 | 1 | | 0.54 | 0.28 | 10 | 1 | | 0.53 | 0.27 | 20 | 1 | | 0.53 | 0.27 |
| | | | 10 | | 0.51 | 0.25 | | 2 | | 0.52 | 0.26 | | 2 | | 0.53 | 0.27 | | 2 | | 0.53 | 0.26 |
| | | | 20 | | 0.52 | 0.26 | | 4 | | 0.53 | 0.27 | | 3 | | 0.53 | 0.27 | | 3 | | 0.52 | 0.26 |
| | | | 30 | | 0.50 | 0.24 | | 6 | | 0.52 | 0.26 | | 4 | | 0.52 | 0.26 | | 4 | | 0.53 | 0.27 |
| | | | 40 | | 0.49 | 0.23 | | 8 | | 0.52 | 0.26 | | 5 | | 0.52 | 0.26 | | 5 | | 0.45 | 0.19 |
| | | | 50 | | 0.48 | 0.21 | | 10 | | 0.52 | 0.25 | | 6 | | 0.52 | 0.26 | | | | | |
| | | | 60 | | 0.46 | 0.20 | | 12 | | 0.52 | 0.25 | | 7 | | 0.53 | 0.27 | | | | | |
| | | | 70 | | 0.43 | 0.17 | | 14 | | 0.52 | 0.26 | | 8 | | 0.51 | 0.24 | | | | | |
| | | | 80 | | 0.33 | 0.10 | | 16 | | 0.47 | 0.21 | | 9 | | 0.45 | 0.19 | | | | | |
| | | | 90 | | 0.20 | 0.03 | | 18 | | 0.38 | 0.13 | | 10 | | 0.29 | 0.72 | | | | | |
| | | | 100 | | 0.01 | -0.01 | | 20 | | 0.12 | 0.00 | | | | | | | | | | |
| HCPD | Sex | 1 | 1 | 0.79 | | | 5 | 1 | 0.78 | | | 10 | 1 | 0.78 | | | 20 | 1 | 0.77 | | |
| | | | 10 | 0.76 | | | | 2 | 0.77 | | | | 2 | 0.77 | | | | 2 | 0.77 | | |
| | | | 20 | 0.75 | | | | 4 | 0.76 | | | | 3 | 0.77 | | | | 3 | 0.76 | | |
| | | | 30 | 0.75 | | | | 6 | 0.77 | | | | 4 | 0.77 | | | | 4 | 0.75 | | |
| | | | 40 | 0.74 | | | | 8 | 0.76 | | | | 5 | 0.76 | | | | 5 | 0.73 | | |
| | | | 50 | 0.73 | | | | 10 | 0.76 | | | | 6 | 0.75 | | | | | | | |
| | | | 60 | 0.70 | | | | 12 | 0.74 | | | | 7 | 0.74 | | | | | | | |
| | | | 70 | 0.68 | | | | 14 | 0.73 | | | | 8 | 0.74 | | | | | | | |
| | | | 80 | 0.65 | | | | 16 | 0.72 | | | | 9 | 0.73 | | | | | | | |
| | | | 90 | 0.59 | | | | 18 | 0.68 | | | | 10 | 0.63 | | | | | | | |
| | | | 100 | 0.44 | | | | 20 | 0.54 | | | | | | | | | | | | |
| | Age | 1 | 1 | | 0.70 | 0.47 | 5 | 1 | | 0.71 | 0.48 | 10 | 1 | | 0.71 | 0.48 | 20 | 1 | | 0.71 | 0.49 |
| | | | 10 | | 0.69 | 0.46 | | 2 | | 0.70 | 0.47 | | 2 | | 0.71 | 0.48 | | 2 | | 0.69 | 0.46 |
| | | | 20 | | 0.68 | 0.44 | | 4 | | 0.70 | 0.47 | | 3 | | 0.69 | 0.46 | | 3 | | 0.68 | 0.44 |
| | | | 30 | | 0.67 | 0.43 | | 6 | | 0.69 | 0.45 | | 4 | | 0.69 | 0.45 | | 4 | | 0.67 | 0.42 |
| | | | 40 | | 0.64 | 0.38 | | 8 | | 0.68 | 0.44 | | 5 | | 0.67 | 0.42 | | 5 | | 0.66 | 0.41 |
| | | | 50 | | 0.62 | 0.36 | | 10 | | 0.66 | 0.42 | | 6 | | 0.68 | 0.44 | | | | | |
| | | | 60 | | 0.60 | 0.34 | | 12 | | 0.67 | 0.43 | | 7 | | 0.66 | 0.42 | | | | | |
| | | | 70 | | 0.54 | 0.26 | | 14 | | 0.64 | 0.39 | | 8 | | 0.65 | 0.40 | | | | | |
| | | | 80 | | 0.46 | 0.18 | | 16 | | 0.62 | 0.36 | | 9 | | 0.65 | 0.40 | | | | | |
| | | | 90 | | 0.33 | 0.08 | | 18 | | 0.58 | 0.32 | | 10 | | 0.47 | 0.20 | | | | | |
| | | | 100 | | 0.02 | -0.03 | | 20 | | 0.21 | 0.01 | | | | | | | | | | |
| PNC | Sex | 1 | 1 | 0.77 | | | 5 | 1 | 0.77 | | | 10 | 1 | 0.76 | | | 20 | 1 | 0.75 | | |
| | | | 10 | 0.75 | | | | 2 | 0.75 | | | | 2 | 0.74 | | | | 2 | 0.74 | | |
| | | | 20 | 0.72 | | | | 4 | 0.73 | | | | 3 | 0.73 | | | | 3 | 0.73 | | |
| | | | 30 | 0.72 | | | | 6 | 0.73 | | | | 4 | 0.73 | | | | 4 | 0.73 | | |
| | | | 40 | 0.71 | | | | 8 | 0.73 | | | | 5 | 0.73 | | | | 5 | 0.69 | | |
| | | | 50 | 0.70 | | | | 10 | 0.73 | | | | 6 | 0.73 | | | | | | | |
| | | | 60 | 0.68 | | | | 12 | 0.72 | | | | 7 | 0.72 | | | | | | | |
| | | | 70 | 0.66 | | | | 14 | 0.71 | | | | 8 | 0.71 | | | | | | | |
| | | | 80 | 0.62 | | | | 16 | 0.69 | | | | 9 | 0.69 | | | | | | | |
| | | | 90 | 0.57 | | | | 18 | 0.65 | | | | 10 | 0.60 | | | | | | | |
| | | | 100 | 0.47 | | | | 20 | 0.54 | | | | | | | | | | | | |
| | Age | 1 | 1 | | 0.54 | 0.28 | 5 | 1 | | 0.53 | 0.27 | 10 | 1 | | 0.53 | 0.27 | 20 | 1 | | 0.53 | 0.27 |
| | | | 10 | | 0.52 | 0.26 | | 2 | | 0.52 | 0.26 | | 2 | | 0.54 | 0.28 | | 2 | | 0.54 | 0.28 |
| | | | 20 | | 0.53 | 0.27 | | 4 | | 0.54 | 0.28 | | 3 | | 0.54 | 0.28 | | 3 | | 0.53 | 0.27 |
| | | | 30 | | 0.52 | 0.26 | | 6 | | 0.53 | 0.27 | | 4 | | 0.53 | 0.27 | | 4 | | 0.52 | 0.26 |
| | | | 40 | | 0.50 | 0.24 | | 8 | | 0.53 | 0.26 | | 5 | | 0.54 | 0.28 | | 5 | | 0.48 | 0.21 |
| | | | 50 | | 0.49 | 0.23 | | 10 | | 0.53 | 0.27 | | 6 | | 0.53 | 0.26 | | | | | |
| | | | 60 | | 0.46 | 0.20 | | 12 | | 0.51 | 0.25 | | 7 | | 0.52 | 0.25 | | | | | |
| | | | 70 | | 0.42 | 0.16 | | 14 | | 0.50 | 0.24 | | 8 | | 0.51 | 0.24 | | | | | |
| | | | 80 | | 0.35 | 0.11 | | 16 | | 0.48 | 0.22 | | 9 | | 0.48 | 0.22 | | | | | |
| | | | 90 | | 0.21 | 0.03 | | 18 | | 0.41 | 0.16 | | 10 | | 0.29 | 0.07 | | | | | |
| | | | 100 | | 0.01 | 0.00 | | 20 | | 0.11 | 0.00 | | | | | | | | | | |

**Table S13. Age and sex predictions across 1%, 5%, 10%, and 20% of features.**

| Phenotype | Decile | Age | | Sex | | Motion | | Minority Representation | | SES | |
|---|---|---|---|---|---|---|---|---|---|---|---|
| | | r | q2 | r | q2 | r | q2 | r | q2 | r | q2 |
| PNC EF | 1 | 0.33 | 0.09 | 0.32 | 0.08 | 0.38 | 0.13 | 0.30 | 0.07 | | |
| | 2 | 0.31 | 0.08 | 0.32 | 0.08 | 0.37 | 0.11 | 0.30 | 0.07 | | |
| | 3 | 0.31 | 0.08 | 0.32 | 0.08 | 0.37 | 0.12 | 0.30 | 0.07 | | |
| | 4 | 0.31 | 0.07 | 0.31 | 0.08 | 0.36 | 0.11 | 0.29 | 0.06 | | |
| | 5 | 0.30 | 0.07 | 0.31 | 0.08 | 0.35 | 0.10 | 0.29 | 0.06 | | |
| | 6 | 0.30 | 0.07 | 0.31 | 0.07 | 0.34 | 0.09 | 0.30 | 0.07 | | |
| | 7 | 0.30 | 0.07 | 0.30 | 0.07 | 0.33 | 0.09 | 0.29 | 0.06 | | |
| | 8 | 0.30 | 0.07 | 0.28 | 0.06 | 0.32 | 0.08 | 0.27 | 0.05 | | |
| | 9 | 0.28 | 0.06 | 0.27 | 0.05 | 0.28 | 0.06 | 0.23 | 0.03 | | |
| | 10 | 0.15 | 0.00 | 0.18 | 0.01 | 0.15 | 0.00 | 0.13 | -0.01 | | |
| PNC Language | 1 | 0.42 | 0.16 | 0.37 | 0.12 | 0.45 | 0.18 | 0.31 | 0.08 | | |
| | 2 | 0.39 | 0.14 | 0.36 | 0.11 | 0.43 | 0.17 | 0.29 | 0.07 | | |
| | 3 | 0.38 | 0.13 | 0.34 | 0.10 | 0.41 | 0.15 | 0.28 | 0.07 | | |
| | 4 | 0.37 | 0.12 | 0.34 | 0.10 | 0.4 | 0.15 | 0.28 | 0.06 | | |
| | 5 | 0.38 | 0.13 | 0.35 | 0.11 | 0.4 | 0.15 | 0.29 | 0.07 | | |
| | 6 | 0.37 | 0.13 | 0.34 | 0.11 | 0.4 | 0.14 | 0.28 | 0.07 | | |
| | 7 | 0.36 | 0.12 | 0.33 | 0.10 | 0.39 | 0.13 | 0.28 | 0.07 | | |
| | 8 | 0.35 | 0.11 | 0.32 | 0.09 | 0.37 | 0.12 | 0.28 | 0.07 | | |
| | 9 | 0.32 | 0.09 | 0.32 | 0.09 | 0.34 | 0.10 | 0.26 | 0.05 | | |
| | 10 | 0.18 | 0.02 | 0.21 | 0.04 | 0.18 | 0.02 | 0.15 | 0.01 | | |
| HCPD EF | 1 | 0.14 | -0.09 | 0.14 | -0.11 | 0.15 | -0.13 | 0.12 | -0.10 | 0.14 | -0.10 |
| | 2 | 0.15 | -0.07 | 0.15 | -0.09 | 0.15 | -0.12 | 0.14 | -0.08 | 0.15 | -0.08 |
| | 3 | 0.15 | -0.07 | 0.15 | -0.08 | 0.15 | -0.11 | 0.14 | -0.08 | 0.15 | -0.08 |
| | 4 | 0.15 | -0.07 | 0.15 | -0.08 | 0.16 | -0.10 | 0.14 | -0.07 | 0.15 | -0.07 |
| | 5 | 0.16 | -0.07 | 0.16 | -0.07 | 0.17 | -0.09 | 0.14 | -0.07 | 0.15 | -0.07 |
| | 6 | 0.16 | -0.07 | 0.16 | -0.07 | 0.17 | -0.08 | 0.14 | -0.07 | 0.15 | -0.07 |
| | 7 | 0.16 | -0.06 | 0.17 | -0.06 | 0.16 | -0.08 | 0.13 | -0.07 | 0.15 | -0.07 |
| | 8 | 0.15 | -0.06 | 0.16 | -0.06 | 0.16 | -0.08 | 0.12 | -0.07 | 0.15 | -0.07 |
| | 9 | 0.15 | -0.05 | 0.14 | -0.06 | 0.13 | -0.08 | 0.10 | -0.07 | 0.14 | -0.06 |
| | 10 | 0.07 | -0.04 | 0.08 | -0.04 | 0.07 | -0.04 | 0.05 | -0.05 | 0.07 | -0.04 |
| HCPD Language | 1 | 0.19 | -0.09 | 0.19 | -0.10 | 0.19 | -0.11 | 0.16 | -0.11 | 0.18 | -0.10 |
| | 2 | 0.15 | -0.13 | 0.15 | -0.14 | 0.15 | -0.14 | 0.12 | -0.14 | 0.13 | -0.14 |
| | 3 | 0.15 | -0.12 | 0.15 | -0.13 | 0.16 | -0.13 | 0.12 | -0.14 | 0.14 | -0.13 |
| | 4 | 0.15 | -0.12 | 0.15 | -0.13 | 0.16 | -0.13 | 0.13 | -0.13 | 0.14 | -0.13 |
| | 5 | 0.16 | -0.11 | 0.16 | -0.11 | 0.16 | -0.13 | 0.14 | -0.12 | 0.15 | -0.11 |
| | 6 | 0.17 | -0.09 | 0.17 | -0.10 | 0.15 | -0.13 | 0.15 | -0.11 | 0.16 | -0.10 |
| | 7 | 0.18 | -0.08 | 0.17 | -0.09 | 0.16 | -0.12 | 0.14 | -0.11 | 0.17 | -0.09 |
| | 8 | 0.17 | -0.08 | 0.16 | -0.09 | 0.15 | -0.11 | 0.13 | -0.11 | 0.16 | -0.09 |
| | 9 | 0.13 | -0.09 | 0.11 | -0.10 | 0.12 | -0.11 | 0.09 | -0.12 | 0.12 | -0.09 |
| | 10 | 0.06 | -0.05 | 0.05 | -0.05 | 0.05 | -0.05 | 0.04 | -0.05 | 0.05 | -0.05 |
| HBN EF | 1 | 0.18 | -0.01 | 0.18 | -0.01 | 0.19 | -0.03 | 0.16 | -0.02 | 0.18 | -0.01 |
| | 2 | 0.15 | -0.03 | 0.15 | -0.03 | 0.16 | -0.05 | 0.13 | -0.03 | 0.15 | -0.03 |
| | 3 | 0.14 | -0.03 | 0.15 | -0.03 | 0.16 | -0.05 | 0.12 | -0.04 | 0.14 | -0.03 |
| | 4 | 0.14 | -0.02 | 0.14 | -0.03 | 0.14 | -0.05 | 0.12 | -0.03 | 0.14 | -0.03 |
| | 5 | 0.14 | -0.02 | 0.14 | -0.03 | 0.13 | -0.05 | 0.12 | -0.03 | 0.14 | -0.02 |
| | 6 | 0.13 | -0.02 | 0.13 | -0.03 | 0.12 | -0.06 | 0.11 | -0.03 | 0.13 | -0.03 |
| | 7 | 0.12 | -0.03 | 0.12 | -0.03 | 0.12 | -0.06 | 0.11 | -0.03 | 0.12 | -0.03 |
| | 8 | 0.11 | -0.03 | 0.11 | -0.03 | 0.10 | -0.06 | 0.11 | -0.03 | 0.11 | -0.03 |
| | 9 | 0.08 | -0.04 | 0.08 | -0.04 | 0.08 | -0.04 | 0.10 | -0.02 | 0.08 | -0.04 |
| | 10 | 0.03 | -0.02 | 0.04 | -0.02 | 0.03 | -0.02 | 0.05 | -0.02 | 0.03 | -0.02 |
| HBN Language | 1 | 0.22 | -0.01 | 0.21 | -0.01 | 0.19 | -0.01 | 0.19 | -0.02 | 0.2 | -0.01 |
| | 2 | 0.21 | -0.01 | 0.20 | -0.01 | 0.18 | -0.01 | 0.18 | -0.02 | 0.19 | -0.01 |
| | 3 | 0.19 | -0.02 | 0.18 | -0.02 | 0.17 | -0.01 | 0.17 | -0.03 | 0.18 | -0.02 |
| | 4 | 0.19 | -0.02 | 0.18 | -0.02 | 0.17 | -0.01 | 0.16 | -0.03 | 0.18 | -0.01 |
| | 5 | 0.19 | -0.02 | 0.19 | -0.01 | 0.17 | -0.01 | 0.17 | -0.02 | 0.18 | -0.01 |
| | 6 | 0.19 | -0.01 | 0.19 | 0.00 | 0.18 | -0.01 | 0.17 | -0.02 | 0.19 | -0.01 |
| | 7 | 0.18 | -0.02 | 0.19 | 0.00 | 0.17 | -0.01 | 0.18 | -0.01 | 0.18 | -0.01 |
| | 8 | 0.16 | -0.02 | 0.18 | -0.01 | 0.16 | -0.01 | 0.17 | -0.01 | 0.17 | -0.01 |
| | 9 | 0.13 | -0.03 | 0.14 | -0.02 | 0.14 | -0.02 | 0.14 | -0.02 | 0.14 | -0.02 |
| | 10 | 0.05 | -0.02 | 0.06 | -0.02 | 0.07 | -0.01 | 0.06 | -0.01 | 0.06 | -0.01 |

**Table S14. Prediction performances controlling for confounds.** Partial correlation was performed at the step in which brain edges were related to each phenotype.

| Phenotype | Percent of Edges | Decile | Performance | |
|---|---|---|---|---|
| | | | r | q2 |
| PNC EF | 10 | 1 | 0.21 | 0.02 |
| | | 2 | 0.22 | 0.02 |
| | | 3 | 0.22 | 0.02 |
| | | 4 | 0.21 | 0.01 |
| | | 5 | 0.20 | 0.01 |
| | | 6 | 0.20 | 0.00 |
| | | 7 | 0.20 | 0.01 |
| | | 8 | 0.20 | 0.01 |
| | | 9 | 0.18 | 0.00 |
| | | 10 | 0.08 | -0.02 |
| PNC Language | | 1 | 0.29 | 0.06 |
| | | 2 | 0.29 | 0.05 |
| | | 3 | 0.28 | 0.05 |
| | | 4 | 0.28 | 0.04 |
| | | 5 | 0.27 | 0.04 |
| | | 6 | 0.27 | 0.04 |
| | | 7 | 0.26 | 0.03 |
| | | 8 | 0.25 | 0.03 |
| | | 9 | 0.22 | 0.02 |
| | | 10 | 0.10 | -0.01 |

**Table S15. PNC rest connectome predictions.** To test the impact of incorporating task fMRI into averaged connectomes on model performances, we repeated PNC analyses using only resting-state connectomes. This analysis was restricted to a subset of participants (n=1063) with available resting-state scans. Model performances across deciles are reported for executive function (top) and language abilities (bottom).
